# Treatment history shapes the evolution of complex carbapenem-resistant phenotypes in *Klebsiella* spp.

**DOI:** 10.1101/2022.08.30.505938

**Authors:** Natalia C. Rosas, Jonathan J. Wilksch, Jake Barber, Jiahui Li, Yanan Wang, Zhewei Sun, Andrea Rocker, Chaille Webb, Laura Perlaza-Jiménez, Christopher Stubenrauch, Dhanasekaran Vijaykrishna, Jiangning Song, George Taiaroa, Mark Davies, Richard A. Strugnell, Qiyu Bao, Tieli Zhou, Michael J. McDonald, Trevor Lithgow

**Affiliations:** Centre to Impact AMR, Monash University, Clayton 3800, Australia; Infection Program, Biomedicine Discovery Institute and Department of Microbiology, Monash University, Clayton 3800, Australia; School of Biological Sciences, Monash University, Clayton 3800, Australia; The First Affiliated Hospital of Wenzhou Medical University, Wenzhou, China; Infection Program, Biomedicine Discovery Institute and Department of Biochemistry & Molecular Biology, Monash University, Clayton 3800, Australia; Wenzhou Medical University, Wenzhou, China; School of Public Health, LKS Faculty of Medicine, The University of Hong Kong, Hong Kong Special Administrative Region, China; Department of Microbiology and Immunology, The Peter Doherty Institute, The University of Melbourne, Parkville 3052, Australia

**Keywords:** antimicrobial resistance, porin, drug-resistance, fitness

## Abstract

Antibiotic resistance is driven by selection, but how a bacterial strain’s evolutionary history shapes drug-resistance remains an open question. Here we reconstruct the genetic and evolutionary mechanisms of carbapenem resistance in a clinical isolate of *Klebsiella*. A combination of short and long read sequencing, machine learning, genetic and enzymatic analyses established that this carbapenem-resistant strain carries no carbapenemase-encoding genes. Genetic reconstruction of the resistance phenotype confirmed that two distinct genetic loci are necessary for the strain to acquire carbapenem resistance. Experimental evolution of the carbapenem-resistant strains in growth conditions without the antibiotic revealed that both loci confer a significant cost, and are readily lost by *de novo* mutation resulting in the rapid evolution of a carbapenem-sensitive phenotype. Thus, historical contingency - a patient’s treatment history - can shape the evolution of antibiotic resistance and suggests that the strategic combinations of antibiotics could direct the evolution of low-fitness, drug-resistant genotypes.

## INTRODUCTION

Understanding the genetic and evolutionary provenance of antimicrobial resistance (AMR) will be important for developing strategies that slow or prevent the evolution of untreatable pathogens. In the case of bacterial pathogens, there is mounting evidence that the source of antibiotic resistance can be the patient’s own microbiome^1^, and that treatment history, as well as pathogen genotype, should be taken into account when designing a treatment plan. Carbapenems are a class of β-lactam antibiotics typically reserved for high-risk, multidrug-resistant infections^2,3^. Surveillance studies show that carbapenem-resistant Enterobacteriaceae (CRE), especially various species of the pathogen *Klebsiella*, have become a widespread problem in clinical settings around the globe^4-6^.

There are two general mechanisms of carbapenem resistance. The first and readily diagnosable mechanism is the acquisition of a single gene encoding a carbapenemase enzyme that directly inactivates the antibiotic. The second mechanism requires multiple genetic loci and bacterial strains in this category often have an over expressed β-lactamase and/or an overexpressed efflux pump and/or an inactivated outer membrane protein^3,7,8^ and potentially as yet unidentified loci as well. Of these two mechanisms, most carbapenem-resistant *Klebsiella pneumoniae* that have been identified so far are caused by carbapenemases. Like other β-lactamases, carbapenemases can be recognized by sequence analysis ^9^ and there are two structurally distinct classes of carbapenemases: the first is the metallo-β-lactamases a family of enzymes that contains a metal ion (usually zinc) coordinated in the active site, a classic example of which is NDM-1^10^. The second carbapenemase family does not coordinate a metal ion but instead relies on an active site serine to hydrolyse carbapenem drugs^7,11^. This second class of carbapenemase includes the IMI, OXA-48 and GES enzymes in addition to the archetypal *<u>K</u>lebsiella <u>p</u>neumoniae* <u>c</u>arbapenemases: the KPC enzymes^11-13^ that include the prevalent KPC-2. The KPC-2 carbapenemase has become widespread, found in many clinical investigations of CRE, and the gene encoding this enzyme - *bla*_KPC-2_ - is transferred readily by horizontal gene transfer via plasmids^14-18^. Since this is a monogenic phenotype, carbapenem-resistance caused by the presence of a carbapenemase is readily diagnosed by whole genome sequencing or even simple PCR-based genome tests.

It has recently become apparent that numerous CRE infections do not depend on the expression of carbapenemases, and there is a mounting evidence that these non-carbapenemase CRE are widespread^19^. Based on the limited cases identified for non-carbapenemase CRE so far, it has been suggested that they emerge as a result of reduced outer membrane permeability and/or increased drug efflux^6,20,21^. Other studies have suggested that, at least in some genetic backgrounds, an extended-spectrum β-lactamase (ESBL) could provide sufficient activity against carbapenems as to generate a CRE phenotype ^7,22^. Despite this emerging knowledge, few studies have directly demonstrated the cause of non-carbapenemase CRE, and the evolutionary forces that shape the evolution of this trait have not been addressed. Given the omnigenic nature of this type of AMR phenotype, understanding the details of these evolutionary forces could provide a means to re-sensitize populations of bacteria to carbapenems.

A patient died as a result of septicaemia caused by a *Klebsiella* isolate, FK688, where the infection did not respond to treatment with carbapenems^23^. In this study (Fig. 1A), we present the complete genome sequence of *Klebsiella* FK688, using a compilation of short-read and long-read sequence data. We identify, and experimentally confirm the genetic basis of the non-carbapenemase CRE phenotype in FK688: carriage of a mega-plasmid (pNAR1) and an inactivating mutation in the chromosomal gene *ompK36*. We show that non-carbapenemase CRE strains are unfit and readily evolve to be drug-sensitive in the absence of carbapenem antibiotics. The evolution of these low-fitness CRE strains may be contingent on their recent exposure to antibiotics that select for other β-lactamases and shows how careful diagnosis and use of antibiotic combinations can drive wanted or unwanted evolutionary outcomes.

**Figure 1.**
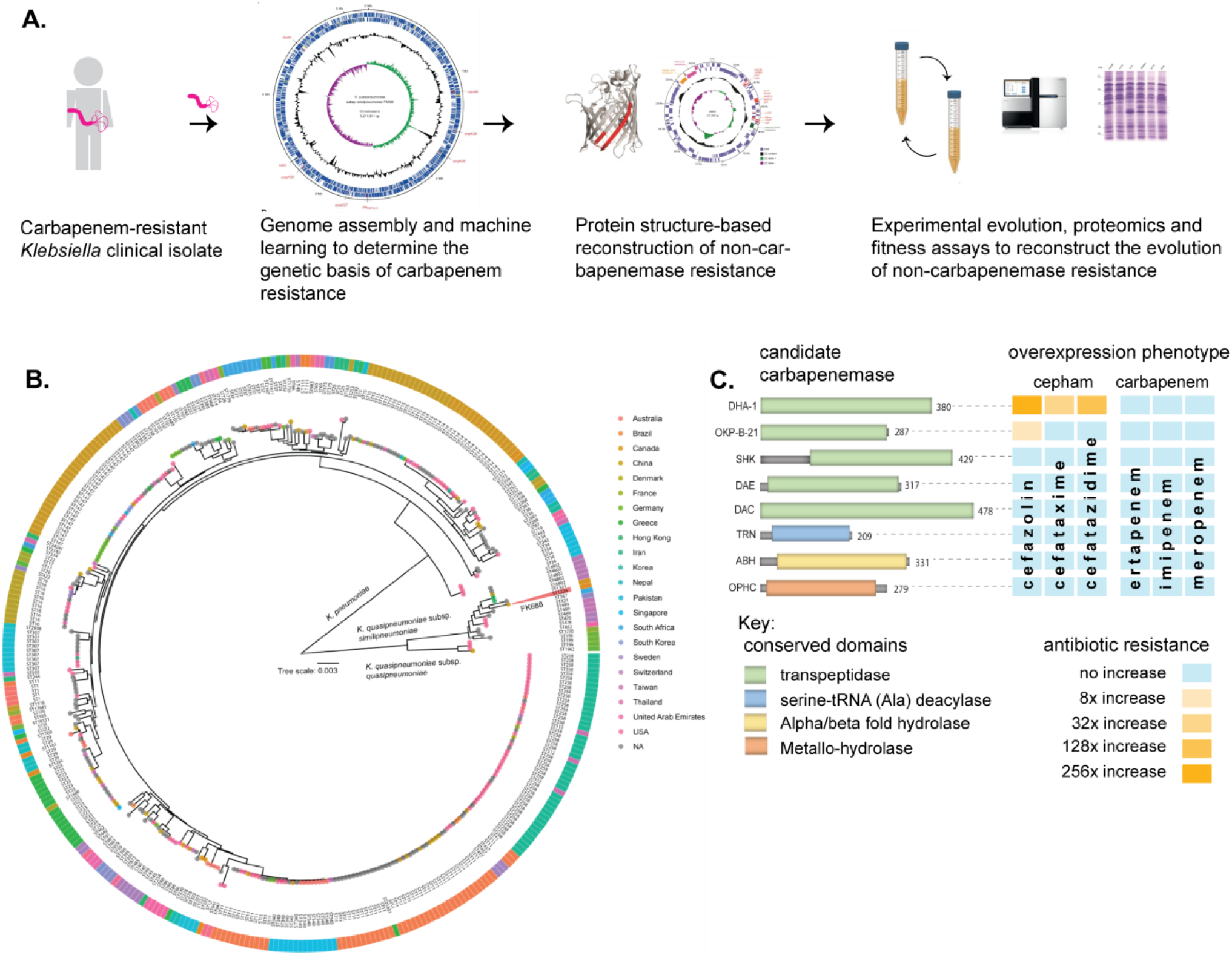
Experiment Overview. (**A**) Carbapenem resistant *Klebsiella spp*. was isolated from a patient and the genome sequenced and assembled. The genetic cause of resistance was confirmed by reengineering the carbapenem resistance, partly based on structure guided restoration of a partially truncated membrane protein. The evolutionary drivers of resistance and sensitivity were determined using experimental evolution and extenstive phenotypic and genotypic measures of evolutionary change. (**B**) The phylogeny analysis of *Klebsiella*. Maximum likelihood phylogenetic tree of 597 publicly available *Klebsiella* genomes showing *K. pneumoniae* and *K. quasipneumoniae* as distinct species. The inner ring indicates the country of strain isolation, as specified in the colour key (NA indicates the genomes with location not deposited in the NCBI). The outer ring colours indicates the distribution of the sequence type classifications. The position of strain FK688 is labelled. (**C**) Eight candidate carbapenemases were identified in the FK688 genome sequence and overexpressed in an *E. coli* model of resistance. Only two genes (DHA1 and OKP-B-21) conferred resistance to the three cepham antibiotics tested, and none of the genes conferred resistance to any carbapenem antibiotic.

## RESULTS

### *Klebsiella* FK688 does not encode a carbapenemase

First, to determine the genetic basis of carbapenem resistance in FK688 (Table 1), we used a long and short-read sequencing approach to generate a complete assembly of the FK688 genome. The genome assembly (On-line Methods) revealed a circular chromosome (5,211,811 bp) and a novel circular megaplasmid (pNAR; 1,257,585 bp). This plasmid carried many of the antibiotic resistance genes corresponding to the known resistance profile^23^, as well as genes encoding efflux pumps and other transporters (Table S1, Table S2). Phylogenetic analysis placed FK688 within *K. quasipneumoniae* subsp. *similipneumoniae* (Fig. 1). Readily identifiable determinants of AMR on the FK688 chromosome are a *bla*_OKP-B-21_ gene encoding a β-lactamase that confers resistance to penicillins and cephalosporins such as cefazolin, as well as determinants for quinolone (*oqxA, oqxB*) ^24-26^) and fosfomycin (*fosA5*) resistance (Fig. S1A).

**Table 1.** Antimicrobial susceptibility profiling of *K. quasipneumoniae* FK688.

| Antimicrobial class | Antimicrobial drug | MIC (µg/mL) <sup>a</sup> |  |  |
| --- | --- | --- | --- | --- |
|  |  | FK688 | *E. coli* (ATCC 25922) | Breakpoints<sup>b</sup> |
| Penicillins | Ampicillin | >2048 | 8 | ≥ 32 |
| Cephems | Cefazolin | >2048 | 2 | ≥ 8 |
|  | Cefotaxime | 1024 | 0.125 | ≥ 4 |
|  | Ceftazidime | >2048 | 0.5 | ≥ 16 |
| Carbapenems | Ertapenem | 64 | 0.016 | ≥ 2 |
|  | Imipenem | 8 | 0.25 | ≥ 4 |
|  | Meropenem | 4 | 0.03 | ≥ 4 |
| Lipopeptides | Polymyxin B | 4 | 2 | ≥ 4 |
| Aminoglycosides | Gentamicin | 1 | 2 | ≥ 16 |
|  | Tobramycin | 1 | 2 | ≥ 16 |
|  | Kanamycin | 2 | 4 | ≥ 64 |
| Tetracyclines | Tetracycline | 128 | 1 | ≥ 16 |
| Fluoroquinolones | Ciprofloxacin | 1 | 0.016 | ≥ 1 |

<sup>a</sup> Drug-sensitive, black text; drug-resistant, red-text
<sup>b</sup> Resistant clinical breakpoint for Enterobacterales given by CLSI<sup>66</sup>

Our genome sequence analysis did not reveal a gene encoding KPC-2, the carbapenemase found in many *K. quasipneumoniae* isolates^25^. To identify any cryptic carbapenemases that may have escaped annotation, we made use of the machine learning predictor DeepBL^9^. DeepBL identifies genes encoding β-lactamases of all types, including carbapenemases, and generated eight high confidence predictions. The two highest predictions represent known beta-lactamases *bla*_OKP-B-21_ and *bla*_DHA-1._ The next closest predictions were for proteins annotated as “serine hydrolase” (SHK), “MBL fold-metallo hydrolase” (OPHC) and “_D_-alanyl-_D_-alanine endopeptidase” (DAE) with conserved domain architecture (CDART) predictions suggesting that our search was sufficiently broad to identify plausible candidates (Fig. 1B, Table S3). To test for carbapenemase activity, we expressed each of the eight genes in *E. coli* and measured growth (Fig. S2; Table S4). MIC assays of *E. coli* expressing the DeepBL candidates showed that only OKP-B-21 and DHA-1 have β-lactamase activity that includes significant resistance to ceftazidime, a third-generation cephalosporin (Fig. 1C, Table S5). However, neither OKP-B-21 nor DHA-1 – nor any of the other proteins tested - provided resistance to carbapenems. Taken together with the genome sequence analysis, these data support that the observed carbapenem-resistant phenotype for FK688 is not caused by a carbapenemase.

### The major porins permit carbapenem sensitivity in *K. quasipneumoniae*

Porins are β-barrel proteins that transport nutrients across the outer membrane of Gram-negative bacteria but can also admit antibiotics into the bacterial cell^27-29^. *Klebsiella pneumoniae* has four genes encoding the major porins OmpK35, OmpK36, OmpK37 and OmpK38^29^, and the position and synteny of each gene in FK688 is conserved across *K. quasipneumoniae* (Fig. S1B, Fig. S3). Inspection of the predicted protein sequences encoded by the four genes in FK688 revealed a 1.3 kb transposase gene (IS*4* family) insertion within the 5’ end of the *ompK35* gene (Fig. S1B), and a 48 bp in-frame deletion in *ompK36* in FK688 (Fig S1B).

The structure of OmpK36 (PDB 5O79) is known^30^, and the identified deletion of 16 amino acids at the 3’ end of the *ompK36* gene in FK688 encompasses large portions of the β14 and β15 strands of the β-barrel structure (Fig. 2A, Fig. 2B) explaining why the protein is not assembled into the outer membrane of FK688. To confirm that this mutation was contributing to carbapenem resistance to we used the *ompK36* gene from *K. quasipneumoniae* subsp. *similipneumoniae* ATCC 700603^31^ to carry out structure-informed repair of the OmpK36 gene in *K. quasipneumoniae* FK688 (Fig. 2C). The resultant strain (*ompK36*^+^pNAR1) was subjected to immunoblotting with antisera, confirming restoration of OmpK36 expression in the *ompK36*^*+*^pNAR1 strain (Fig. 2D). Finally, measurements of carbapenem MIC determined that the repaired gene encoding the porin OmpK36 restored carbapenem sensitivity (Table 2).

**Table 2.** Antimicrobial susceptibility profiling of FK688-derived strains.

| Antimicrobial class | Antimicrobial drug | MIC (µg/mL) <sup>a</sup> |  |  |  |  |
| --- | --- | --- | --- | --- | --- | --- |
|  |  | *ΔompK36* |  |  | *ompK36*<sup>+</sup> |  |
|  |  | pNAR1 | pNAR1<br>*Δbla<sub>DHA-1</sub>* | pNAR1<sup>−</sup> | pNAR1 | pNAR1<br>*Δbla<sub>DHA-1</sub>* |
| Penicillins | Ampicillin | > 2048 | 256 | 128 | > 2048 | 128 |
| Cephems | Cefazolin | > 2048 | 32 | 32 | > 2048 | 4 |
|  | Cefotaxime | 1024 | 0.5 | 0.5 | 32 | 0.125 |
|  | Ceftazidime | > 2048 | 1 | 0.5 | 512 | 0.25 |
| Carbapenems | Ertapenem | 64 | 0.5 | 0.5 | 0.5 | 0.016 |
|  | Imipenem | 8 | 0.25 | 0.25 | 1 | 0.12 |
|  | Meropenem | 4 | 0.06 | 0.125 | 0.125 | 0.03 |
| Lipopeptides | Polymyxin B | 4 | 4 | 4 | 4 | 4 |
| Aminoglycosides | Kanamycin | 2 | 4 | 2 | 4 | 4 |
| Tetracyclines | Tetracycline | 128 | 256 | 2 | 128 | 128 |
| Fluoroquinolones | Ciprofloxacin | 1 | 0.06 | 0.06 | 1 | 0.125 |

<sup>a</sup> Drug-sensitive, black text; drug-resistant, red-text

**Figure 2.**
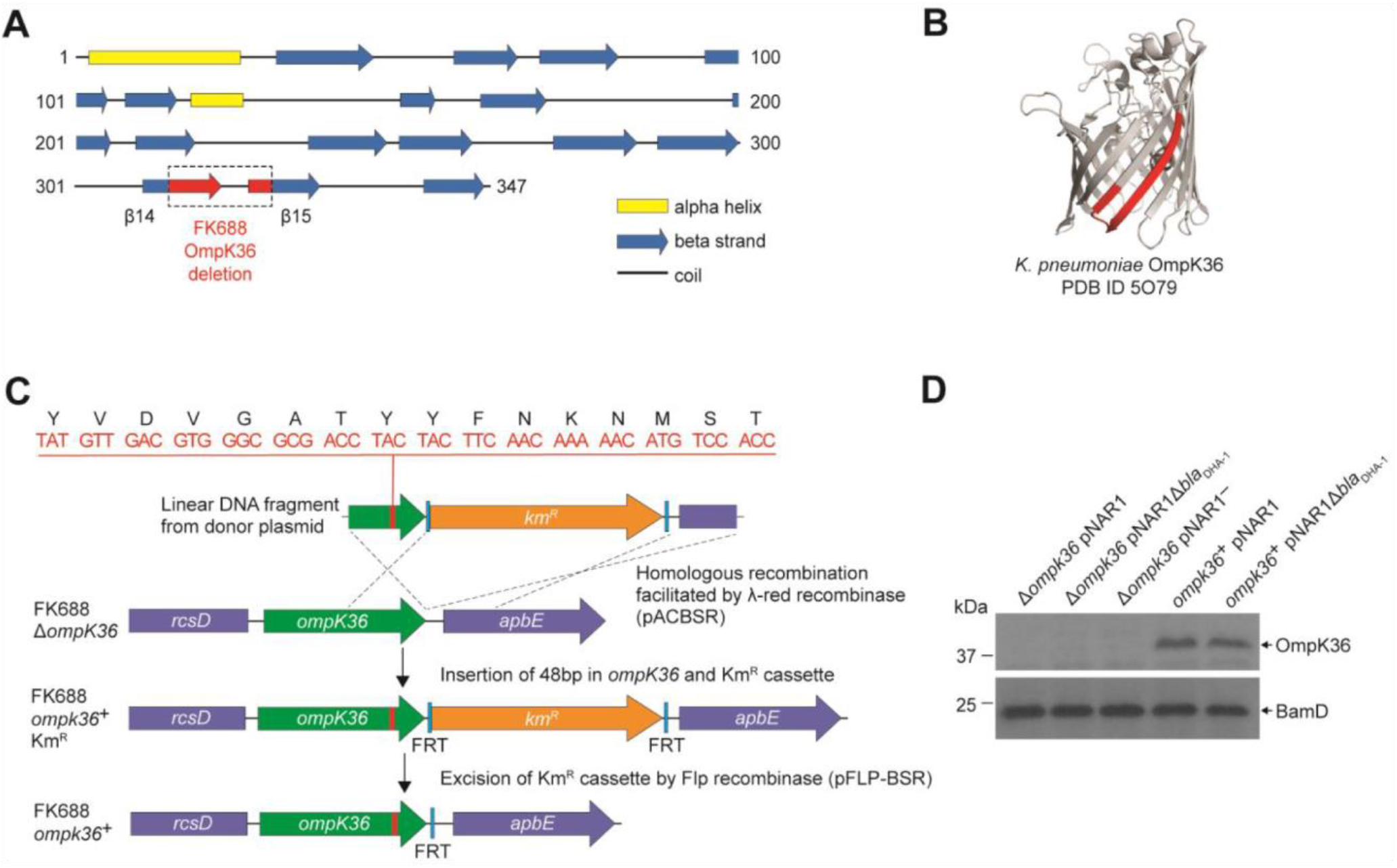
Reconstruction of mutant OmpK36 protein based structural characteristics of OmpK36 in *K. quasipneumoniae* subsp. *similipneumoniae*. (A) We repaired the mutant OmpK36 by identifying the missing DNA sequence based on the PSIPRED^81^ secondary structure prediction of the OmpK36, using the protein sequence encoded in the *K. quasipneumoniae* subsp. *similipneumoniae* genome. The location of the 16 amino acid deleted region in the FK688 OmpK36 is highlighted in red on the β14-β15 strands of the structural model. (B) Tertiary structure prediction for the β-barrel OmpK36 monomer (PDB ID 5O79^30^). (C) Schematic depicting the engineering approach used to restore a functional version of *ompK36* in FK688 by the insertion of 48 nucleotides in *ompK36*, as shown. FRT (flippase recognition target) sites permitted excision of the Km^R^ (kanamycin resistance) cassette using Flp recombinase. Following Km^R^ excision, a single FRT site and scar region remain in between the *ompK36* and *apbE* genes. The amino acid identity between the OmpK36 from ATCC 700603 and FK688 is 95% (Supplementary Fig. S2B) and the ATCC 700603 sequence^31^ was used to repair the *ompK36* locus of FK688, as described in the Methods section. (D) Total membrane extracts were prepared from the indicated strains, the proteins in the samples analyzed by SDS-PAGE and immunoblotting using an antibody probe that recognizes OmpK36 ^29^. The outer-membrane protein BamD was used as a sample loading control for the analysis.

### DHA-1 and Δ*ompK36* are required for carbapenem resistance and impose non-additive fitness costs in growth media without antibiotics

To determine the evolutionary stability of the pNAR1 plasmid, we passaged 10 replicate mutation accumulation lines of *K. quasipneumoniae* FK688 in growth media without β-lactam antibiotic selection (Fig. 3A). After 11 passages, two replicates had completely lost resistance to the β-lactam antibiotic ceftazidime. The first lineage lost a 17 kb region of pNAR1 that included the *bla*_DHA-1_ and *qnrB4* antibiotic resistance genes flanked by the gene mobility elements *tnpA*-*sul1* (pNAR1Δ*bla*_DHA-1_) while the second lineage lost the entire pNAR1 plasmid (pNAR1‾) (Fig. 3B, 3C). We assayed both the pNAR1‾ and pNAR1Δ*bla*_DHA-1_ for growth and antibiotic sensitivity and confirmed that loss of pNAR1, and specifically the *tnpA*-*sul1* region of pNAR1, caused a loss of resistance to ceftazidime and carbapenem (Table 2, Fig. S4).

**Figure 3.**
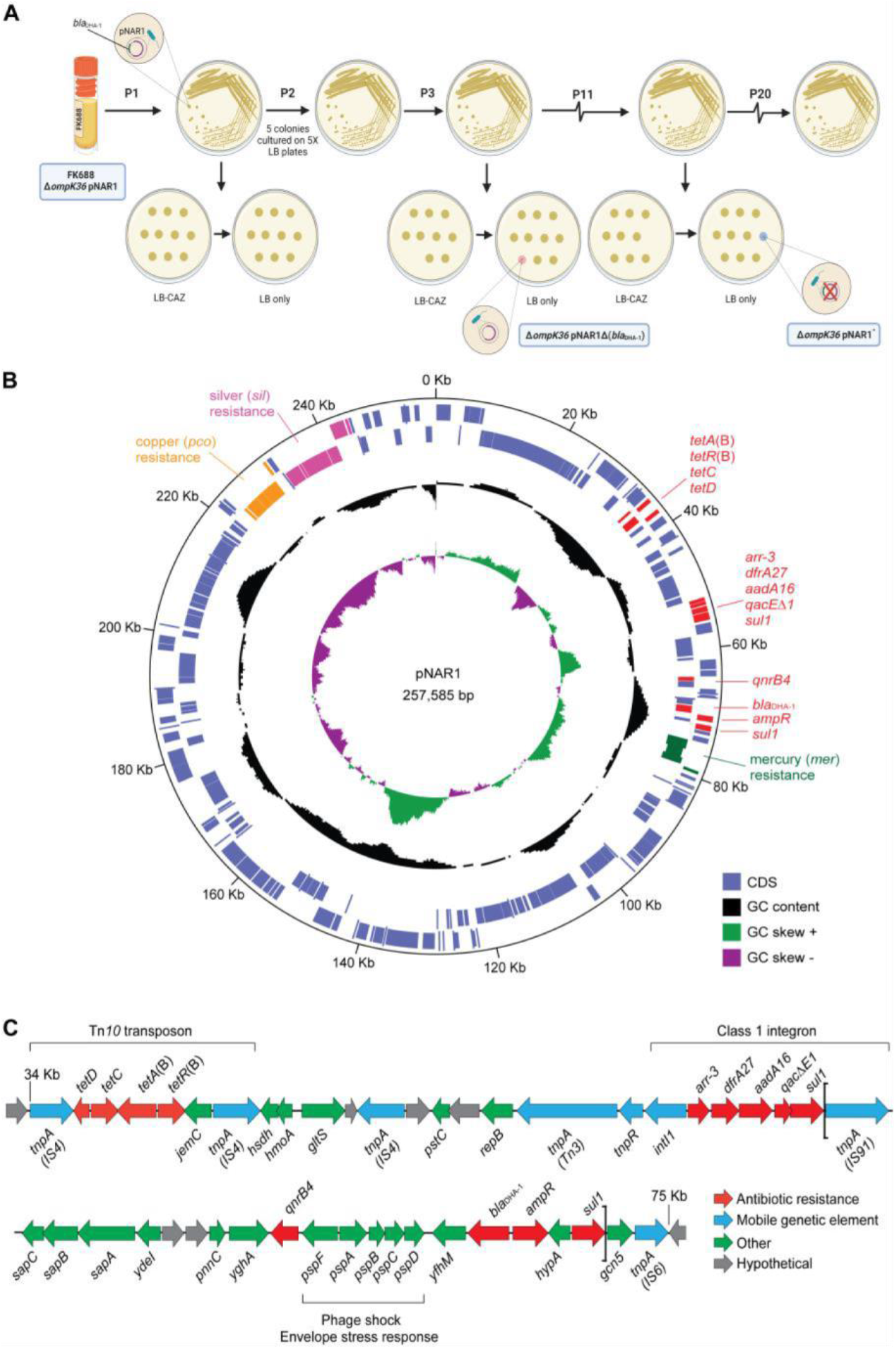
Evolution and physical map of plasmid pNAR1. (A) Schematic representation of the *in vitro* evolution experiment. After passage #3 (P3) a ceftazidime-susceptible (CAZ^S^) mutant evolved, lacking a 17 kb region of pNAR1 that included *bla*_DHA-1_ (referred to as Δ*ompK36* pNAR1Δ*bla*_DHA-1_). After 11 passages (P11) a CAZ^S^ colony missing the entire plasmid (referred to as Δ*ompK36* pNAR1^−^) evolved. In total, 20 passages were performed, and another five CAZ^S^ colonies were identified, each missing the 17 kb region of pNAR1 that includes *bla*_DHA-1_. (B) The position of genes encoding antibiotic resistance determinants (red), and efflux pumps annotated as being for mercury resistance (green), copper resistance (orange), and silver resistance (pink) are indicated. In addition to *bla*_DHA-1_, pNAR1 carries genes encoding AmpR (ID00077) a transcriptional regulator known to regulate expression of *bla*_DHA-182_. Also, other drug resistance genes including those responsible for resistance to tetracycline (*tetA*(B)), rifamycin (*arr-3*), trimethoprim (*dfrA27*), streptomycin (*aadA16*), macrolides (*qacΔE1*), sulfonamides (*sul1*), and quinolones and fluoroquinolones (*qnrB4*) (Supplementary Fig. S5, Supplementary Table S1). The blue lines in the outer concentric circles represent the location of predicted coding sequences in the forward (outer most) and reverse DNA strands. The middle circle (black) indicates the % GC content, and the inner circle indicates the positive (green) and negative (purple) GC skew [(G-C)/(G+C)]. The map was generated with DNAPlotter^79^. (C) Linear map of a 41 kbp segment of pNAR1 showing the genetic arrangement of antimicrobial resistance genes (red), mobile genetic elements (blue), annotated coding sequences (green), and hypothetical genes of unknown function (grey). Assigned IS families are shown underneath each transposase gene (*tnpA*). The loci within the two brackets represents the 17 kbp DNA segment (*tnpA*-*sul1*) deleted from pNAR1Δ*bla*_DHA-1_

Whole genome sequencing established that the pNAR1‾ and pNAR1Δ*bla*_DHA-1_ strains had not sustained any other mutations, which confirmed the genotypes of strains Δ*ompK36* and *ompK36*^+^ with and without the plasmid-encoded *bla*_DHA-1_ gene. We therefore engineered repaired versions *ompK36*^+^ for each strain and tested each mutant for growth and carbapenem resistance (Table 2). The results demonstrate that the acquisition of carbapenem-resistance in FK688 required the combination of (i) the absence of porins, and (ii) the β-lactamase DHA-1 (Table S6).

While the expression of DHA-1 and the absence of a functional porin provide a selective advantage to *Klebsiella* in high concentrations of carbapenem antibiotics, we sought to understand whether these genotypes impose a fitness cost in environments without carbapenems. Competitive fitness assays were carried out to determine the fitness effects of the *bla*_DHA-1_ and *ompK36* alleles alone and in combination (Methods, Fig. 4A). We found that the *bla*_DHA-1_ and *ΔompK36* alleles conferred a substantial fitness cost in growth media without antibiotic as compared to the other strains tested (Fig. 4B). To refine our understanding in terms of carbapenem-resistance, we measured the fitness effects of the *bla*_DHA-1_ and *ompK36* alleles across a range of imipenem concentrations. These assays showed that the fitness defects seen in the absence of imipenem (Fig. 4B) are gradually reversed in the strains grown in the presence of increasing concentrations imipenem (Fig. 4C). At the breakpoint value of 0.125 µg /mL, three genotypes are already at a selective disadvantage: *ompK36*^+^pNAR1Δ*bla*_DHA-1_, *ompK36*^+^pNAR1 and Δ*ompK36* pNAR1‾. Above the breakpoint value at 0.25 µg/mL, all four genotypes are at a selective disadvantage relative to the parental FK688. Thus, a clinically-relevant appearance of the CRE phenotype requires a combination of the presence of DHA-1 and the lack of a functional porin.

**Figure 4.**
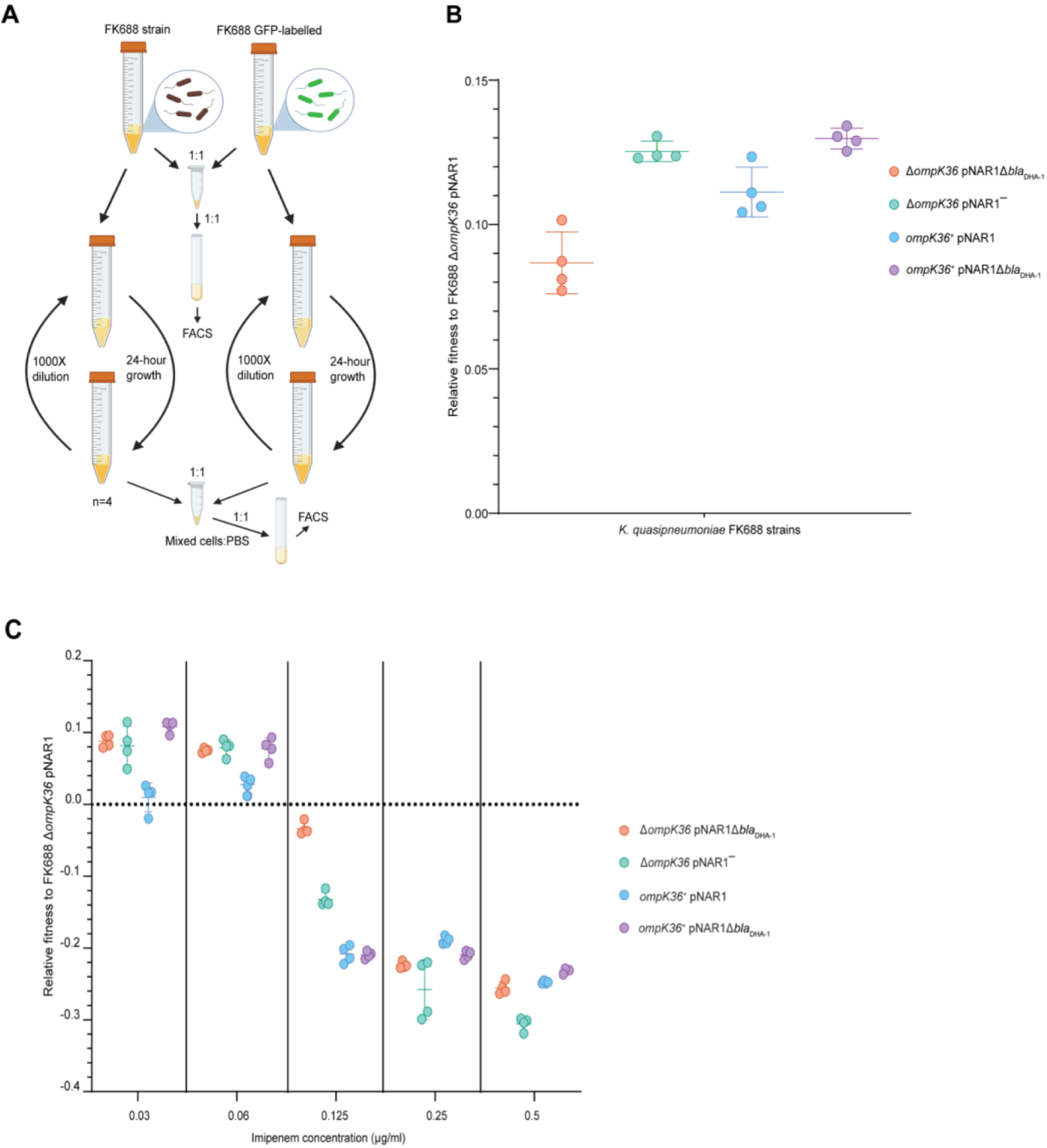
Competitive fitness assay of FK688 strain variants against GFP-labelled FK688. (A) Schematic of the competitive fitness assay experiment (Methods). FACS: fluorescence-activated cell sorting. (B) The relative fitness of the engineered mutant strains relative to the carbapenem resistant FK688 strain, measured in LB growth media without antibiotics. Mutant strains either have the OmpK36 outer membrane transporter restored (*ompK36+*), the DHA1 beta-lactamase deleted (pNAR1- or pNAR1Δ*bla*_*DHA-1*_), or both (purple circles). The y-axis shows the selection coefficient (S) per generation compared to the carbapenem resistant ancestor FK688 which has its fitness set at 0. The legend indicates the genotypes for each strain (note FK688 is genotype Δ*ompK36* pNAR1). Error bars represent mean with SD. (C) Relative fitness of FK688 mutant strains compared to parental FK688, measured in LB media supplemented increasing concentrations of imipenem. The legend shows the genotype for each *Klebsiella* strain. Error bars represent mean ±SD.

### High fitness and carbapenemase sensitivity rapidly evolves in experimental populations of FK68

Mutations that inactivate major porins restrict the permeability of the outer membrane. To address the fitness cost of major porin loss over time, we passaged 20 Δ*ompK36* and 20 *ompK36*^+^ replicate populations across 200 generations of evolution in media without antibiotics (Fig. 5A). The Δ*ompK36* populations evolved similar competitive fitness to the *ompK36*^+^ evolved populations, recovering the fitness cost of the Δ*ompK36* mutation (Fig. 5A). MIC analysis showed that the Δ*ompK36* populations had evolved increased sensitivity to imipenem, 64-fold for the two FK688 Δ*ompK36* populations that we tested (Table 3). Analysis of the whole genome sequence data for four clones found that they had each sustained a deletion in pNAR1 that removed *bla*_DHA-1_. This explains the increased carbapenem sensitivity of the Δ*ompK36* lineages as they had now lost one of the genes contributing to provide the phenotype.

**Table 3.** **Antimicrobial susceptibility profiling FK688 Δ*ompK36* and *ompK36*^*+*^ strains and their respective evolved strains.** ^a^Drug-sensitive, black text; drug-resistant, red-text

| Antimicrobial class | Antimicrobial drug | MIC ($\mu\text{g/mL}$) <sup>a</sup> | | | | | |
| --- | --- | --- | --- | --- | --- | --- | --- |
| | | $\Delta ompK36$ | | | $ompK36^+$ | | |
| | | pNAR1 | pNAR1 $\Delta bla_{DHA-1}$ A2(o) | pNAR1 $\Delta bla_{DHA-1}$ A2(t) | pNAR1 | pNAR1 $\Delta bla_{DHA-1}$ B3(o) | pNAR1 $\Delta bla_{DHA-1}$ B3(t) |
| Cephems | Cefazolin | >2048 | 32 | 1 | >2048 | 2 | 1 |
|  | Cefotaxime | 1024 | 0.5 | 0.25 | 32 | 0.125 | 0.25 |
|  | Ceftazidime | >2048 | 0.5 | 0.25 | 512 | 0.25 | 0.25 |
| Carbapenems | Ertapenem | 64 | 0.5 | 0.031 | 0.5 | 0.016 | 0.031 |
|  | Imipenem | 8 | 0.125 | 0.125 | 1 | 0.25 | 0.125 |
|  | Meropenem | 4 | 0.125 | 0.016 | 0.12 | 0.016 | 0.016 |
| Tetracyclines | Tetracycline | 128 | 128 | 128 | 128 | 128 | 128 |
| Fluoroquinolones | Ciprofloxacin | 1 | 0.125 | 0.25 | 1 | 0.031 | 0.063 |
| Aminoglycosides | Kanamycin | 2 | 2 | 2 | 4 | 2 | 2 |
|  | Tobramycin | 0.5 | 1 | 1 | 1 | 1 | 1 |
|  | Gentamicin | 0.5 | 1 | 0.5 | 0.5 | 0.5 | 0.5 |
| Lipopeptides | Polymyxin B | 4 | 4 | 4 | 4 | 8 | 4 |

**Figure 5.**
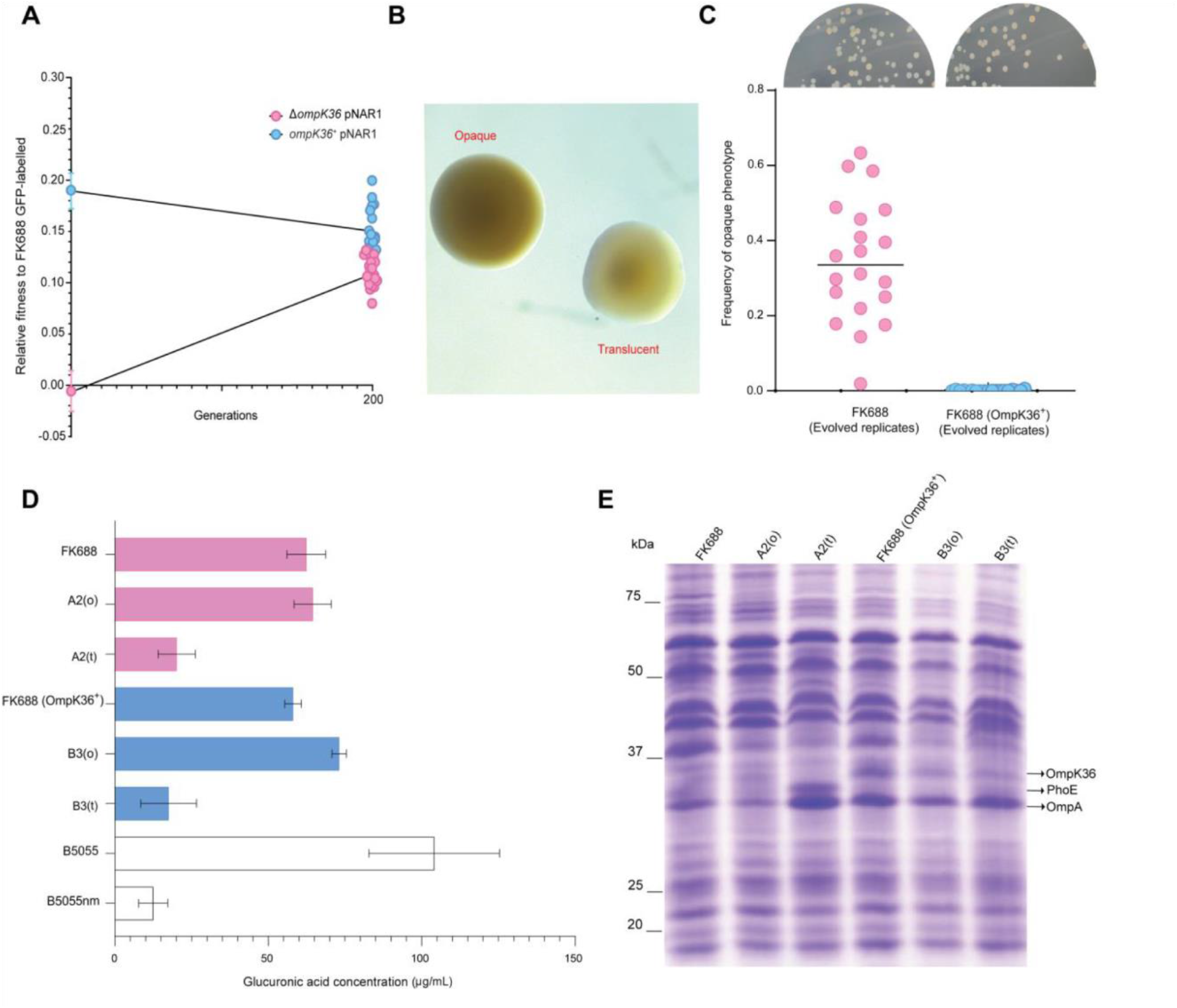
Genotypic and phenotypic evolution of FK688 Δ*ompK36* and *ompK36*^*+*^ strains. (A) The relative fitness assessments for the FK688 Δ*ompK36* (pink) and *ompK36*^*+*^ (blue) genotypes. The error bars represent mean ±SD (n=4). Relative fitness assays were also performed for 20 evolved populations after 200 generations of evolution in LB growth media without antibiotics (right). The line represents individual replicates with means connected. (B) Colony morphotypes seen in the evolved Δ*ompK36* (pink, denoted A2) *ompK36*^*+*^ (blue, denoted B3) populations with a representative opaque (o subscript) and translucent (t subscript) colony indicated. Colonies were cultured on 0.5X LB agar after overnight incubation at 37°C and photographed with stereo microscope using transmitted light. Each dot represents an individually evolved population (C) Relative numbers of opaque colonies in the replicate populations of FK688 (Δ*ompK36*) and (*ompK36*^*+*^) strains after 200 generations. (D) Capsular polysaccharide was extracted from cell cultures for glucuronic acid measurement (methods) ^71^.The error bars represent mean ±SD (n=3). (E) Total cell extracts from the indicated strains were analyzed by SDS-PAGE and Coomassie staining. The migration positions of OmpK36, PhoE and OmpA are indicated. The identities of these protein species were confirmed by mass spectrometry of the corresponding region of the gel.

The ancestral FK688 strain has an opaque colony morphology. After the 200 generation evolution experiment, samples were plated on agar and were found to display a mixture of opaque and translucent colony morphotypes (Fig. 5A, Fig. 5B). This feature has been seen before in *Klebsiella* spp. and depends on the expression of type 1 fimbriae^32 33,34^. The FK699 Δ*ompK36* population evolved to become a mixture of opaque and translucent colony forming types, while in the ompK36^+^ population, the population was entirely made up of translucent colony forming cells (Fig. 5C). In the B3 population a transposase gene (IS*4* family) insertion had disrupted the downstream region of the *fimE* gene (Fig. S5B, Table S7). With only the *fimB* gene left intact, we hypothesize that the B3 population remains translucent because all cells are fimbriated. Fimbrial expression impacts on capsular polysaccharide production: non-fimbriated clones appear mucoid due to capsule secretion and look opaque, while translucent colonies are fimbriated and non-mucoid^32,35,36^. The *fimE* gene was not disrupted in the A2 population that was capable of phase-switching (Fig. S5A). Consistent with these observations, an assay measuring glucuronic acid that reflects the presence of capsule showed that the A2(o) and B3(o) had more capsular polysaccharide than A2(t) and B3(t) (Fig. 6D).

**Figure 6.**
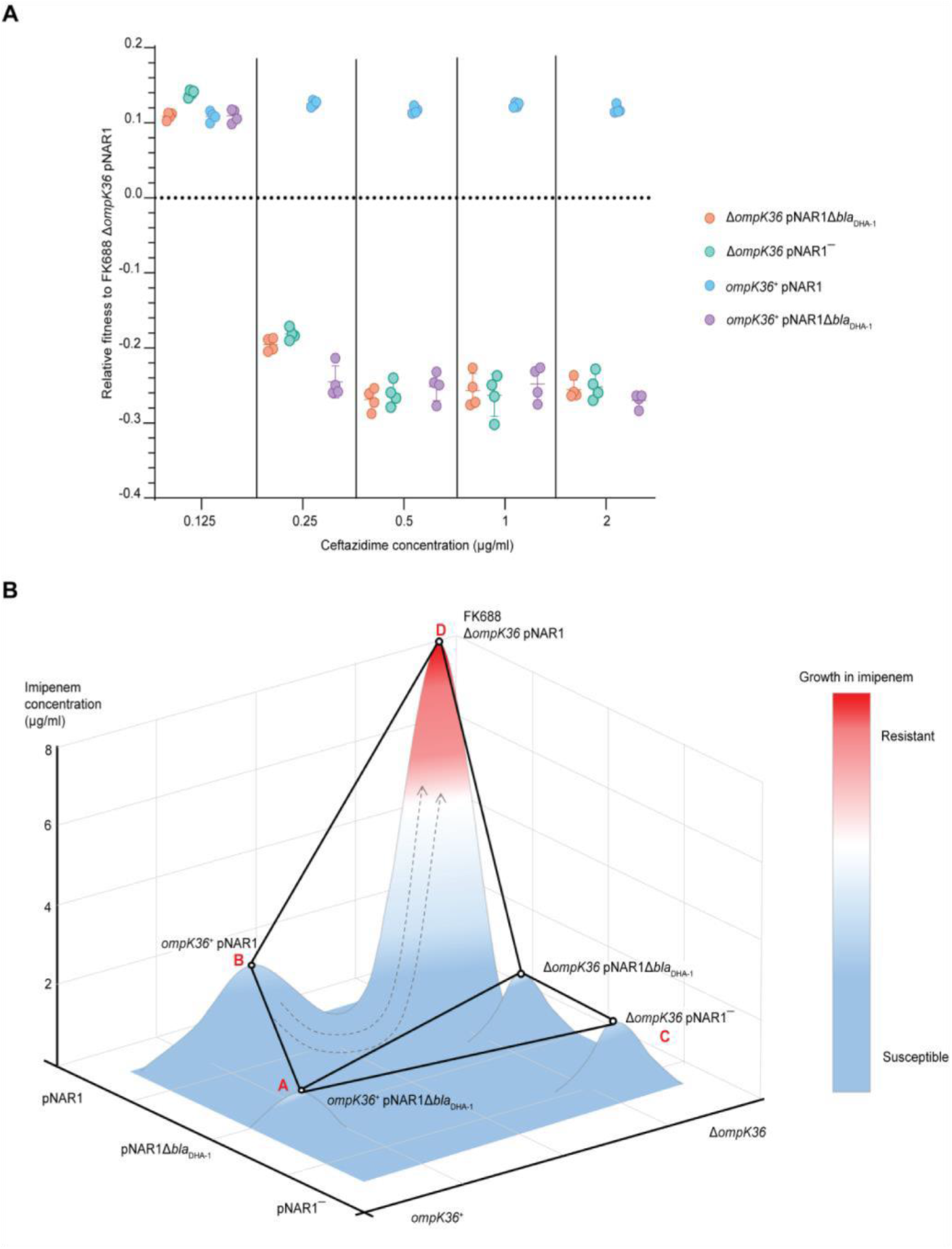
Competitive fitness assay of FK688 strain variants in the presence of ceftazidime. (A) The fitness of FK688 mutants measured in LB media supplemented with ceftazidime across a concentration range from 0.125 to 2 µg/mL. Only strains with an intact pNAR1 plasmid, including the BLA_DHA-1_ gene are able to survive high concentrations of ceftazidime. The legend has the genotypes for the *Klebsiella* strains. Error bars represent mean with SD. (B) Schematic of the imipenem resistance landscape. Each genotype is depicted as being resistant (red) or susceptible (blue) to imipenem. The x and y planes depict the AMR genotypes, and the z plane represents growth measured at each concentration of imipenem. Circles represent the genotype of each strain and lines show strain connected by a single mutation. The evolution of imipenem resistance requires two genes - the *bla*_DHA-1_ gene, and a loss of function mutation in *ompK36*: these two alleles are both found in FK688, indicated at “D”. Since both single-step mutants “B” and “C” are imipenem susceptible, and also low fitness in growth media without drugs (Fig. 4B), we propose that the population had recently been exposed to conditions that selected for the pNAR1 plasmid. Then after exposure of the population to imipenem, the Δ*ompK36* mutant was strongly selected. This suggests that the most likely evolutionary path to imipenem resistance was A – B – D.

We found no differences in fitness or antibiotic resistance for the opaque and translucent types. The crucial outcome of the evolution experiment was the observation that all 20 of the A (i.e. Δ*ompK36*) replicate populations had become more sensitive to imipenem, with evolution of drug-sensitivity being of great interest. Protein analysis by SDS-PAGE and mass spectrometry (Fig. 6E) showed no changes that were consistent in both A2(o) and A2(t) to explain the increased sensitivity to imipenem. Our reconstruction experiments (Table 2) suggest that the primary determinant of this reversion of the antimicrobial resistance phenotype is the observed loss of the *bla*_DHA-1_ gene from the megaplasmid pNAR1.

### Non-carbapenemase carbapenem resistance evolves via ceftazidime resistance

The results so far confirm that two genetic variants are required for the evolution of non-carbapenemase resistance to imipenem. The evolution of this trait is puzzling because both the Δ*ompK36* strain and the *bla*_DHA-1_ positive strain each has a low fitness in growth media without antibiotic, as well as in growth media with imipenem. One explanation for the evolution of carbapenem resistance in a strain of *Klebsiella* with an intact major porin is the simultaneous acquisition of multiple genetic variations after the population was exposed to imipenem. However, given that the two genes are not linked, the simultaneous acquisition of a new gene (*bla*_DHA-1_) and a spontaneous genetic variant (inactivating mutation in one or more major porins such as *ompK36*) is highly unlikely.

An alternative explanation is that the bacterial population may have first been subjected to environmental conditions that selected for the fixation of one of the alleles. For antibiotic resistance phenotypes, this is a realistic scenario. Given the difficulty with diagnosing a non-carbapenemase CRE infection, patients might first be treated with other β-lactams. To address this possibility, we tested whether an antibiotic other than imipenem could have selected for the presence of one or both alleles, thus potentiating the evolution of imipenem resistance with a single mutational step.

The fitness of each combination of the *bla*_DHA-1_ and *ompK36* alleles were tested in a range of ceftazidime concentrations (Fig. 7A). This showed that *bla*_DHA-1_ was selectively favoured, even at concentrations of ceftazidime approximately 100× below the clinical breakpoint of 16 µg/mL despite the fact that carriage of the *bla*_DHA-1_ gene bears a substantial fitness cost (Fig. 5B). In addition, we found that the OmpK36 inactivating mutation reduced fitness in ceftazidime, confirming that selection on ceftazidime can select for the *bla*_DHA-1_ gene, but not for the *ompK36* mutation, which is required for imipenem resistance.

These results support the hypothesis that imipenem resistance evolved in FK688 via multiple evolutionary steps. First, the population experienced an antibiotic-containing environment that selected for the *bla*_DHA-1_ gene. This could have been any antibiotic that selected for the pNAR1 plasmid, such as ceftazidime or another β-lactam antibiotic. After a short period of selection on this first treatment, most of the population would carry the *bla*_DHA-1_ gene, increasing the chance that an *ompK36* inactivating mutation would occur in a cell that also carried the *bla*_DHA-1_ gene. We tested the evolutionary potential for each genotype to evolve imipenem resistance by setting up cultures of the FK688 strains with either one, or none, of the two drug resistance alleles and plating 10^9^ cells of each genotype on a range of concentrations of imipenem (Fig. 6B). We evaluated the evolutionary path to antibiotic resistance by considering the starting point of evolution as the genotype that does not have the *bla*_DHA-1_ gene or the loss of function mutation in *ompK36*, indicated at “A”. This genotype is the most logical starting point because it has the highest fitness in growth conditions without antibiotic (Fig. 5B). We found that strains that carried only the Δ*ompK36* allele, or neither allele, were unable to evolve imipenem resistance (Fig. 6B; “A” and “C”). However, a *K. quasipneumoniae* strain with the pNAR1 plasmid (carrying the *bla*_DHA-1_ gene) would be readily able to evolve resistance (Fig. 7B; “B”). Thus, in a scenario representing previous treatment of a patient with ceftazidime, given the low fitness benefit conferred by the loss of function mutation in *ompK36* in growth media supplemented with imipenem, the path A→B→D is most likely.

## DISCUSSION

This study presented the first physical genetic map of a *K. quasipneumoniae* subsp. *similipneumoniae* genome. In addition to species-specific AMR loci on the bacterial chromosome, this multidrug-resistant strain carries many characterized AMR traits on a megaplasmid that was named pNAR1. We have further used whole genome sequencing of *K. quasipneumoniae* populations to characterise the genetic and evolutionary mechanism with which they acquire carbapenem resistance. A major finding of the study is the ease with which carbapenem sensitivity was restored in the absence of drug selection. This bodes well for new strategies that are being devised to reverse the evolution of AMR phenotypes in *Klebsiella* spp. populations, be they in built environments, in gut microbiomes or in infection sites. A further major finding is the means by which an ill-chosen drug treatment, for example with ceftazidime, can prime a population of *K. quasipneumoniae* to rapidly evolve a carbapenem-resistant (CRE) phenotype.

Genomics-based surveillance has shown that KPC-2 carbapenemases are widespread in *Klebsiella* spp. including *K. quasipneumoniae*^25,37^. β-lactamases encoded by chromosomal genes are common amongst these species of *Klebsiella*: *bla*_SHV_ is found in *K. pneumoniae, bla*_OKP-A_ in *K. quasipneumoniae* subsp. *quasipneumoniae* and *bla*_OKP-B_ in *K. quasipneumoniae* subsp. *similipneumoniae*^26,38,39^. Consistent with this, FK688 carries a chromosomally located *bla*_*OKP-B*._ In addition, FK688 also carries the β-lactamase *bla*_DHA-1_ on the megaplasmid pNAR1.

### Epistatic impacts of porins, pumps and enzymes on carbapenem-resistance

There have been no reports on drug efflux pumps in *K. quasipneumoniae* but in some strains of *K. pneumoniae*, drug-resistance phenotypes have been suggested to include epistatic contributions from genes encoding efflux pumps^28^. FK688 encodes numerous ABC-type efflux systems with annotations for these efflux pumps suggestive of metal ion ligands (copper, silver, mercury), but the ligand specificity of efflux pumps can be broader or different to that denoted by annotation^40,41^. In FK688, the efflux pumps did not contribute to carbapenem resistance. Are there any further epistatic effects relevant to imipenem sensitivity? A potential one would be that other porins had been upregulated, and the A2(t) translucent strain was observed to have increased expression of outer membrane proteins OmpA and PhoE (Fig. 6E). However, (i) OmpA does not form a sizeable channel in the outer membrane^42^, and (ii) while the porin PhoE does form a channel^29^, it is not conducive to permitting imipenem entry into *Klebsiella*^29^. A single SNP that was fixed in the A2 population was in the *cadBA* operon that generates cadaverine (Table S8). CadC is a positive activator of the *cadBA* operon ^43^, and this activation of *cadBA* is known to close porins in general and block β-lactam influx^44-46^, with point mutations in the CadC protein sufficient to inhibit the *cadBA* operon^47^.

Instead, we found that the CRE phenotype in FK688 depends on epistasis between *bla*_DHA-1_ and *ompK36*. Both the carriage of *bla*_DHA-1_ and the defect in *ompK36* have measurable fitness cost to the strain. After multiple rounds of plating in the absence of drug-selection, we observed the loss of a segment (*tnpA-sul1*) of pNAR1 that carries *bla*_DHA-1_; that is, selection against the β-lactamase. This same outcome was also observed in a controlled experimental evolution experiment over 200 generations. These observations are explained by the measured fitness cost in pNAR1Δ*bla*_DHA-1_ which was less than the fitness cost imparted by the full plasmid. Thus, in the absence of β-lactam the CRE phenotype is reversed to carbapenem-sensitivity. It is not clear from current literature how widespread the non-carbapenemase CRE phenotype is, but several points are worthy of note: (i) the presence of plasmids encoding DHA-1 is geographically wide-spread^48^, (ii) the presence of *ompK35* and/or *ompK36* mutations is prevalent in the various species of *Klebsiella*^29^, and (iii) a recent case study showed a single hospital had collected 87 isolates of CRE *Klebsiella* with 55% of them being a non-carbapenemase CRE phenotype^21^.

### Diagnosis and treatment of non-carbapenemase CRE

To obtain the best outcome from the limited treatment options effective against CRE, a personalized approach to antibiotic dosing has been urged^2,49,50^. This in turn requires rapid and accurate diagnosis. All of the currently available tests for CRE aim to identify specific carbapenemases: the Carba NP test detects OXA-48 type carbapenemases, the modified Hodge test identifies metallo-β-lactamases^51^, and the newer more sophisticated gene-specific tests also have limitations^52-56^. The finding that in some environments perhaps half of all CRE cases could be caused by strains that do not encode a carbapenemase^21^, and our finding that DHA-1 expression can - through epistasis with porin mutations - deliver a CRE phenotype, adds a further degree of difficulty to diagnosis of CRE.

While treatment options for CRE are limited^50,57^ this study adds benefit in two important aspects. Firstly, that combination therapy using an ESBL inhibitor such as avibactam or tazobactam could be a good option for this type of CRE^58^. For example, the use of ceftolozane-tazobactam combination therapy is suggested as an alternative to carbapenems in treatment of some CRE infections^59^, and a cohort study of 391 patients with ceftriaxone-resistant infections showed piperacillin-tazobactam combination therapy compared favourably with carbapenem treatment^60^. In strains like FK688, where it is a β-lactamase such as DHA-1 that contributes to carbapenem resistance, specific inhibition of that β-lactamase with avibactam or tazobactam would increase the level of carbapenem or cephalosporin in the bacterial periplasm, and thereby increase the effectiveness of drug treatment. Secondly, our study cautions that moves towards phage therapy being used to treat CRE should not make use of phages that use OmpK35 or OmpK36 as their receptor ^61^. These phages would place selective pressure on the *Klebsiella* strain to become porin-defective, since mutations to inactivate the receptor is a prime cause of phage-resistance^62,63^. Phages that use OmpK35 or OmpK36 as their receptor, would thereby select for porin-defects and thus inadvertently select for carbapenem-resistance; phages used therapeutically should therefore target alternate receptors ^61^.

Finally, this study shows how the evolutionary history of a pathogenic strain can predispose for an AMR phenotype to evolve via specific genetic routes. The likelihood that antibiotic resistance will evolve depends on the strength of selection and the availability of genetic variants that confer resistance to the antibiotic. If the genes or genetic variants that confer resistance to the antibiotic confer a decrease in fitness in environments without antibiotic, then these variants will be exceedingly rare (or absent) from the population. If antibiotic resistance genes are not supplied via horizontal gene transfer, then antibiotic resistance must evolve by the *de novo* mutation of a gene already present in the genome. Since the spontaneous acquisition of multiple genetic variants (for example, Δ*ompK36* and *bla*_DHA-1_) is much less likely than the acquisition of a single new gene (KPC-2 carbapenemase), the evolution of non-carbapenem resistance via two loci seemed unlikely. However, if a strain already carries the *bla*_DHA-1_ gene then the spontaneous evolution of a loss-of-function mutation in an extant chromosomal gene – *ompK36* – is more likely than the spontaneous acquisition of a carbapenemase gene. Exposure to ceftazidime, or any antibiotic that selects for the *bla*_DHA-1_ gene, would therefore potentiate the evolution of carbapenem resistance by a single loss-of-function mutation in major porins. These results show how historical contingency, or an individual’s treatment history, can shape the evolution of AMR, and suggests that the strategic combinations of antibiotics could direct the evolution of low-fitness, antibiotic-resistant genotypes.

## MATERIALS AND METHODS

### Chemicals and reagents

Ampicillin and tetracycline were purchased from Astral Scientific. All other antibiotics were purchased from Sigma-Aldrich in highest possible grade. A stock solution of Anhydrotetracycline (Cayman Chemical Company) in 50% ethanol was prepared to induce β-lactamase production when required.

### Bacterial strains, oligonucleotides, and cultures conditions

Plasmids, bacterial strains and oligonucleotides used in this study are described in Table S4, S9 and S10, respectively. Bacterial cultures were routinely grown in Lysogeny Broth (LB) or cation-adjusted Müller-Hinton Broth (CAMHB) media at 37 ºC with shaking at 200 rpm, unless otherwise stated. When required, antibiotics used for the selection of antibiotic resistance markers were supplemented in growth media at the following concentrations: ampicillin 100 μg/mL; kanamycin 30 μg/mL; chloramphenicol 34 μg/mL; ceftazidime: 0.125 μg/mL, 0.25 μg/mL, 0.5 μg/mL, 1 μg/mL or 2 μg/mL; imipenem: 0.03 μg/mL, 0.06 μg/mL, 0.125 μg/mL, 0.25 μg/mL or 0.5 μg/mL.

### Genome sequencing and evaluation

Genomic DNA of the *Klebsiella* isolates was prepared from solid media scrapings of pure culture using the GenElute Bacterial Genomic DNA Kit (Sigma-Aldrich) and the Gram-negative bacteria protocol. High molecular weight DNA was then isolated using a 0.6× ratio of sample (200 µl) to AMPure XP-beads (120 µl) (A63882, Beckman Coulter). Genomic DNA was sequenced in parallel on the Oxford Nanopore GridION and Illumina Nextseq 500. A detailed explanation of the library preparation and genome sequence annotation and analysis is described in On-line Methods.

### Plasmid annotation

Prokka v1.14.0 was employed to predict pNAR1 genes. The translated gene sequences were used to search against NCBI nr and CARD resistance databases (Comprehensive Antibiotic Resistance Database) with the blastp algorithm and Resistance Gene Identifier (RGI) software (https://github.com/arpcard/rgi), respectively (e value ≤ 10^−5^). The predicted genes received an annotation file containing credible resistance genes (Cut-off as “Perfect” or “Strict”) and putative resistance genes (Cut-off as “Loose”).

### Membrane protein analysis

Total (outer and inner) membranes from *K. quasipneumoniae* were purified following the method of Dunstan et al, 2017^64^ with minor modifications. A complete description is included in On-line Methods.

### Minimum Inhibitory Concentration (MIC) determination

Antimicrobial susceptibility testing was performed by broth microdilution method using CAMHB according to the guidelines the Clinical and Laboratory Standards Institute (CLSI) M07-10th Ed. Document^65^. The resistance of antimicrobial agents was interpreted according to the criteria of CLSI^66^. The assays were performed in biological triplicate with at least two technical replicates. *E. coli* ATCC® 25922 was used as a quality-control strain.

### Mutant construction

A *K. quasipneumoniae* FK688 strain containing a repaired *ompK36* gene (*ompK36*^+^) was constructed using the “gene gorging” technique^67-69^. A donor plasmid was made that contained the repaired *ompK36* gene upstream of a kanamycin-resistant cassette and ∼500 bp of FK688 genomic region downstream of the *ompK36* gene and flaked by I-*Sce*I-endonuclease recognition sites. The PCR products were gel purified, cloned into pJET1.2/blunt and confirmed by sequencing. The primers used are listed in Table S10. The donor plasmid and pACBSR carrying L-arabinose-inducible I-*Sce*I endonuclease and λ-Red recombinase genes, were transformed into FK688 by electroporation. Co-transformants were inoculated into LB containing chloramphenicol and 0.2% (*w/v*) L-arabinose and incubated overnight at 30°C with shaking. Engineered strains were isolated on LB-agar containing kanamycin, cured of the donor and pACBSR plasmids (by their sensitivity to chloramphenicol), and mutants were confirmed by PCR. The self-curing plasmid pFLP-BSR was then used to excise the kanamycin cassette^68^.

### Validation testing of candidate β-lactamases

The coding sequences of β-lactamase and DeepBL candidate genes were amplified from FK688 genomic DNA using Fusion High-Fidelity DNA polymerase (New England BioLabs) with the oligonucleotide primers listed in Table S10. The PCR products and the anhydrotetracycline (ATc)-inducible expression vector pJP-CmR were digested with restriction enzymes using either *Nco*I and *Hin*dIII, or *Eco*RI and *Hin*dIII (New England BioLabs) and ligated to create the plasmids listed in Table S4. Plasmids were verified by sequencing, transformed into *E. coli* BW25113 or *K. quasipneumoniae* FK688 and selected with chloramphenicol. Target gene expression was induced with 35 ng/mL ATc. The parental plasmid pJP-CmR was used as the control in all experiments.

### Plasmid maintenance assessment

A mutation accumulation experiment was performed to evaluate pNAR1 stability in the FK688 Δ*ompK36* pNAR1 strain. The strain was cultured on LB agar (no antibiotics) from a glycerol stock, corresponding to passage #1 (P1). From this plate, 10 colonies were replica cultured on LB agar containing 10 µg/ml ceftazidime (LB-CAZ) and LB agar without antibiotics (LB-only). Five colonies from P1 were individually sub-cultured on five LB agar plates without antibiotics (P2). One colony from each P2 plate was similarly passaged to P20, with replica plating of 10 colonies on LB-CAZ and LB-only after each passage. Therefore, 50 colonies were screened after each passage. Colonies that grew on LB-only but not on LB-CAZ (CAZS) were assessed for pNAR1 maintenance by PCR.

### Fitness assays evaluation

Competitive fitness assays of strains relative to a GFP-expressing reference *K. quasipneumoniae* FK688-GFP^+^ strain were performed as described in Barber *et al*.^70^ with some modifications. Single colonies of ancestral, evolved and reference strains were grown overnight at 37°C in 3 mL LB media (with and without antibiotic selection) with shaking in separate 15 mL falcon tubes. At saturation, the strain of interest and reference strain were mixed (100 μL:100 μL) diluted in PBS and measured by fluorescence-activated cell sorting (FACS) to determine the unadjusted proportions of the two strains. Based off these unadjusted values, volumes of experimental and reference strains were then modified to create a 1:1 cell density ratio, which is the initial starting frequency. The mixture of strains was then diluted 1:1000 (3 μL in 3 mL of LB) before propagating into fresh LB media each day (10 generations per day). 500μL of sample was taken each day, diluted in 1× <u>p</u>hosphate-<u>b</u>uffered <u>s</u>aline (PBS [pH7.4]: 137 mM NaCl, 2.7 mM KCl, 10 mM Na_2_HPO_4_, 2 mM KH_2_PO_4_), and measured by flow cytometry (LSR Fortessa X20a) for the proportion of experimental to reference strains. A maximum total count of 50,000 events was used. The selection coefficient (S) per generation for each experimental strain relative to the reference strain was calculated by taking the natural logarithm of the ratio of experimental to reference strains at the initial and the final time point, and dividing by the number of generations passed^71^ as described by the following regression model formula:

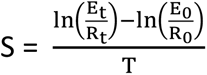

Where T = time (generations); E = frequency of evolved strain; R = frequency of the reference (GFP-labelled) strain; E_t_, R_t_ = frequencies at time “t”; E_0_, R_0_ = initial frequencies.

In other words, when S=0, the strains are equally fit; when S is positive, the evolved strain is more fit than the reference strain; and when S is negative, the evolved strain is less fit than the reference strain.

### Evolution experiments

To set up the evolution experiment, single clones of the drug-resistant, non-functional porin (Δ*ompK36* pNAR1) strain and its homologous, membrane engineered (*ompK36*^*+*^ pNAR1) strain were used to seed twenty replicates each in 3mL LB media, and serially passaged in antibiotic-free media for 24 hours at 37 °C with shaking in separate 15 mL falcon tubes. Every 24 hours the populations were diluted 1:1000 (3 μl in 3 mL of LB), equating to roughly 10 generations per day. The strain populations were evolved for approximately 200 generations. The relative growth rate of all lineages and the parental strains was measured every 50 generations using fitness assays, and colony morphological features were assessed throughout the experiment. After 200 generations, four single clones of evolved strains with different morphological characteristics were selected for whole genome sequencing. After every 20 and 50 generations, 500 μL of evolved strains were mixed with 500 μL of 50% glycerol and stored at -80 °C.

### Growth curves

Overnight bacterial cultures were subcultured 1:100 in CAMHB and grown until mid-log phase (OD_600_ of 0.6-0.8). Cells were then diluted in CAMHB to an OD_600_ of 0.05. Three biological replicates (each in technical replicates) were grown in a 96-well plate using the Tecan Spark 10M. The plate was enclosed in a hydration chamber for 24 hours at 37 °C with orbital shaking (200 rpm) amplitude 3mm, and cell density (OD_600_) measurements were recorded every 60 min.

### Capsule polysaccharide analysis

Capsular polysaccharides of *Klebsiella* strains were extracted by the phenol-extraction method^72^ and quantified using a colorimetric assay for glucuronic acid as previously described^73^. See On-line Methods for details.

### Phylogeny

597 complete *Klebsiella* genomes from the NCBI database were used to make a global phylogeny (downloaded May 2020). Roary (3.11.2)^74^ was used to align the genomes and extract the core genomes. The core genomes were used to generate a phylogenetical tree using RAxML v8.2.12^75^ with a general time reversible nucleotide substitution model with rate heterogeneity modelled with a gamma distribution (GTR+GAMMA). Branch supports were estimated using 1,000 bootstrap replicates. Kleborate v0.3.0^76^ was used to identify MLST types. Visualization of the tree was generated with R using the package ggtree.

## ACKNOWLEDGEMENTS

This work was supported by the National Health & Medical Research Council (grant number 1092262) and a Monash BDI Postgraduate Research Scholarship. Complete genome sequence data for *Klebsiella quasipneumoniae* subsp. s*imilipneumoniae* FK688 has been deposited at the NCBI (Accessions: CP072505, CP072506 and CP072507) and annotated through the Prokaryotic Genome Annotation Pipeline (PGAP) under the Bioproject PRJNA717371 and Biosample SAMN18498882.

