## supplemental figures, tables and methods for "Treatment history shapes the evolution of complex carbapenem-resistant phenotypes in *Klebsiella* spp."

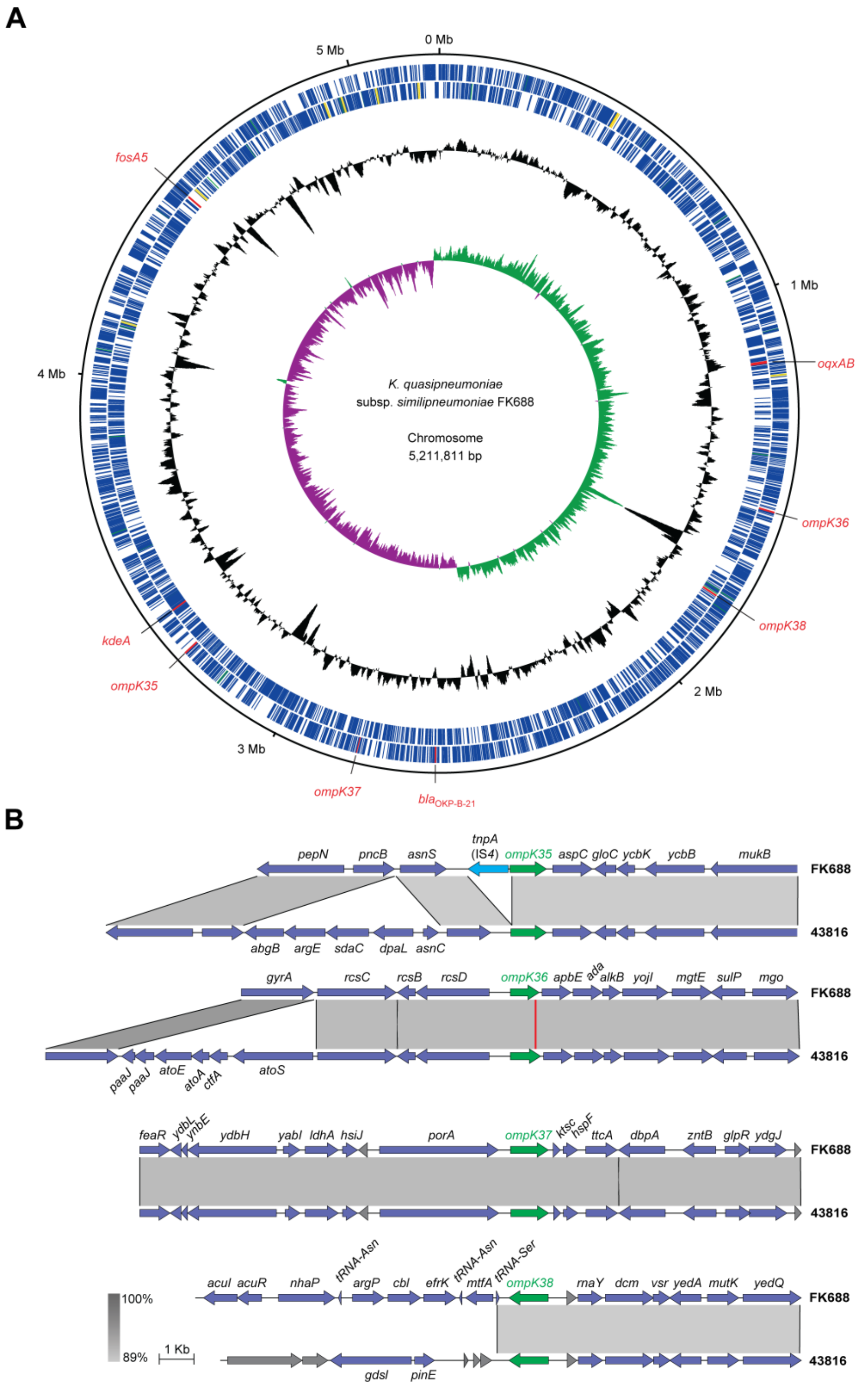


**Figure S1. Physical map of the *K. quasipneumoniae* subsp. *similipneumoniae* FK688 chromosome.** (A) The position of genes encoding antibiotic resistance determinants (*oqxAB*, $bla_{\text{OKP-B-21}}$, *kdeA*, and *fosA5*) and major outer membrane porin proteins (*ompK35*, *ompK36*, *ompK37* and *ompK38*) are indicated. The blue lines in the two outer concentric circles represent the location of predicted coding sequences in the forward (outermost) and reverse DNA strands. The middle circle (black) indicates the % GC content, and the inner circle indicates the positive (green) and negative (purple) GC skew [(G-C)/(G+C)]. The map was generated with DNAPlotter [79]. (B) Gene synteny comparison alignments of *ompK35*, *ompK36*, *ompK37* and *ompK38* (green arrows) and neighbouring ORFs with either predicted (blue arrows) or unknown (grey arrows) functions in FK688 and *K. pneumoniae* subsp. *pneumoniae* ATCC 43816 reference strain. Comparisons were created with BLASTn and the Easyfig application[80] using the default parameters (min. length=0, max. e value=0.001, min. identity value=0). The red line was drawn to indicate the position of a 48-bp intragenic region within *ompK36* present in ATCC 43816 and absent in FK688, which was not detected at the level of resolution used in BLASTn comparative analysis. The nucleotide sequence percentage identity between strains is represented by the gradient indicator (bottom left).

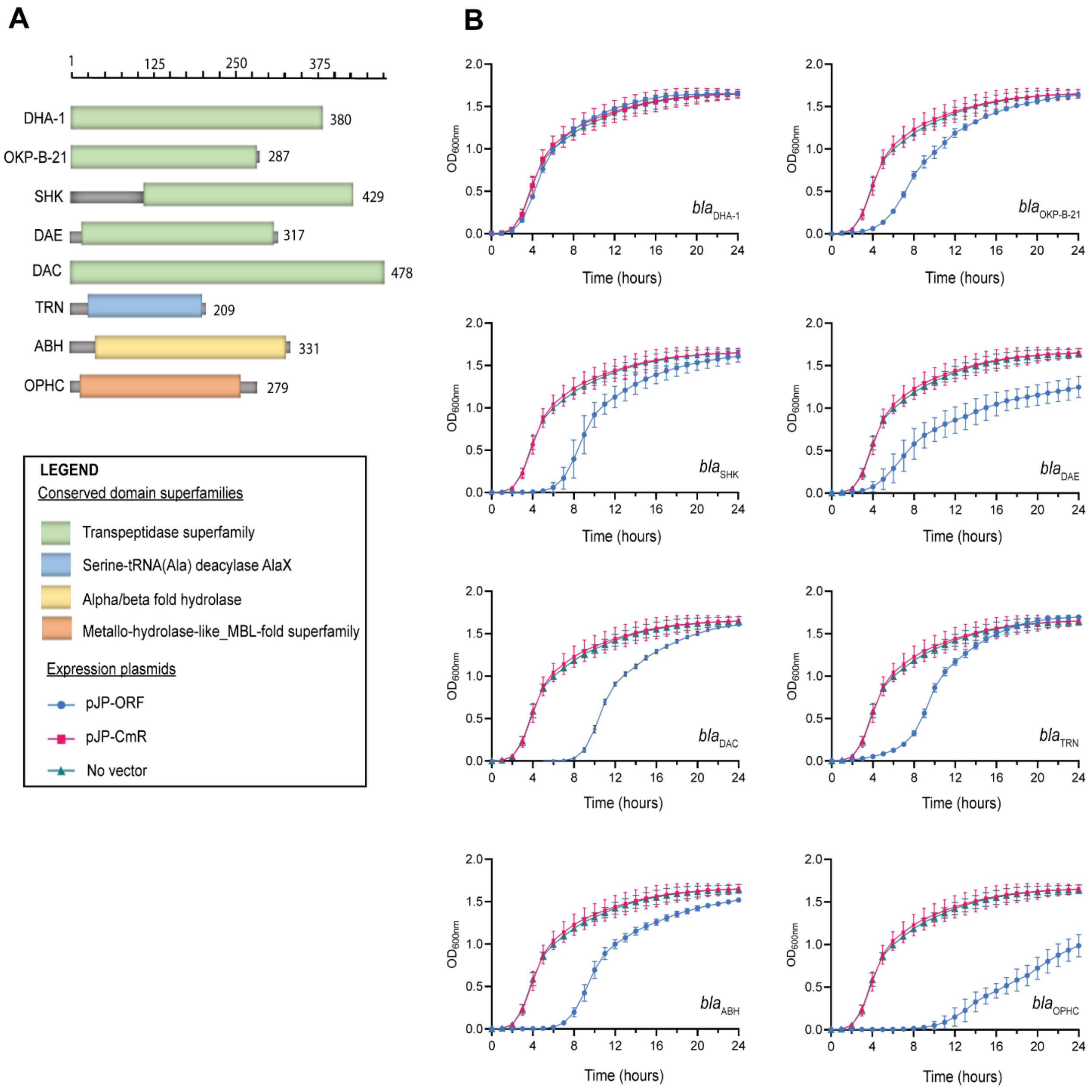


**Figure S2. Domain architecture of DeepBL candidates.** (A) The Conserved Domain Architecture Retrieval Tool (CDART)[83] was used to create the graphical display of domain architectures for the protein sequences identified from DeepBL analysis of the FK688 genome sequence data. DHA-1, OPK-B-21 and several other of the open-reading frames show protein architecture of the Class C β-lactamase (green). TRN, ABH and OPHC have a domain architecture similar to Serine-tRNA deacylase (blue), Alpha/Beta fold hydrolase (yellow) and Metallo-hydrolase-like_MBL-fold superfamily (orange), respectively. Numbers map the amino acid residues for each protein. (B) Growth rate analysis of *E. coli* BW25113 strains expressing the indicated open-reading frames cloned into plasmid pJP-CmR. In each case, the comparison is made between untransformed BW25113 (“no vector”, green triangles), and BW25113 transformed with the parental plasmid (“pJP-CmR”, red squares) or the plasmid carrying the indicated open-reading frame (blue dots). Strains were cultured in LB for 24 hours at 37°C and cell density ($OD_{600}$) was measured every hour. Error bars represent the standard deviation of biological triplicates.

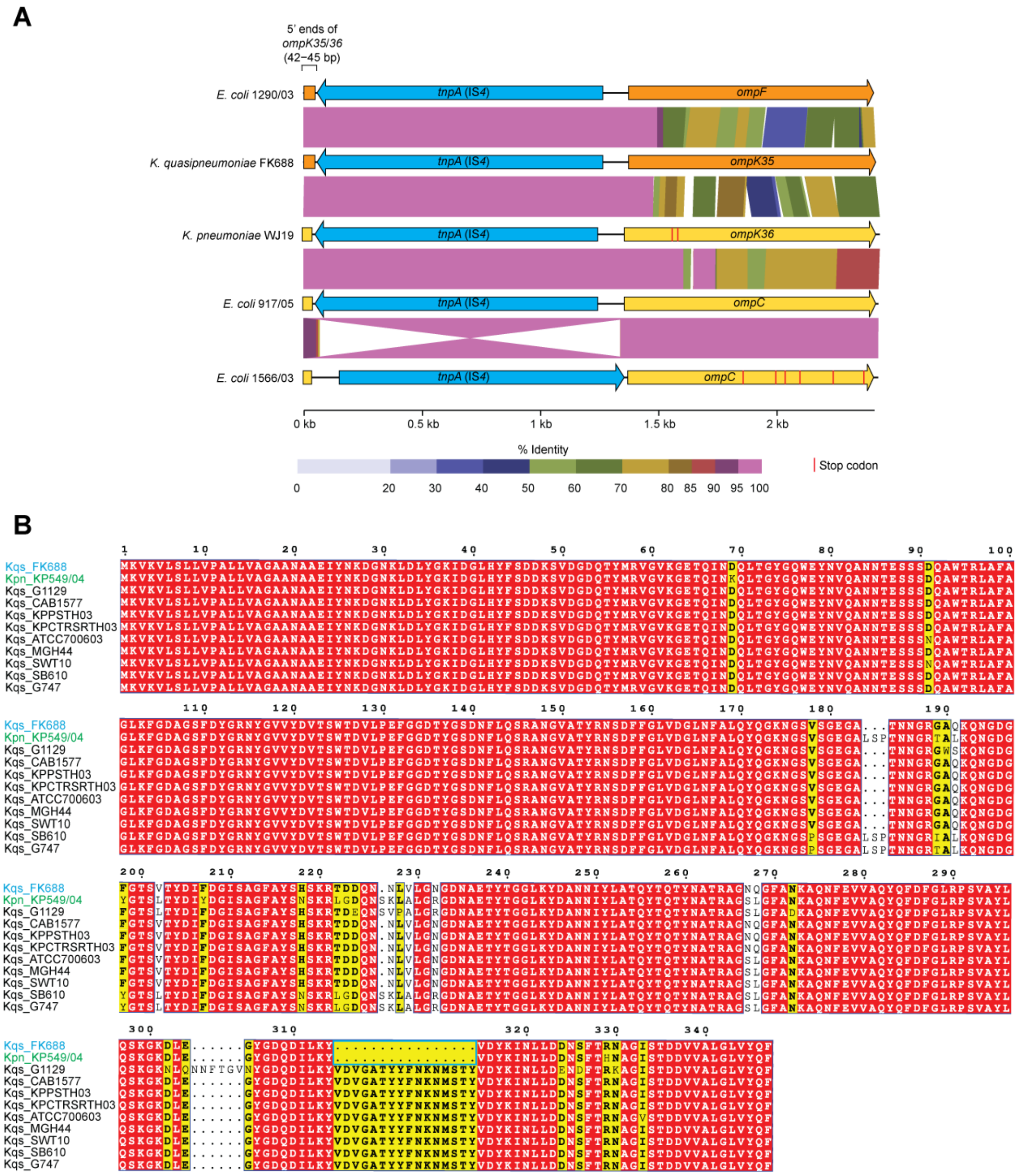


**Figure S3. Comparative sequence analysis of *ompK35* and *ompK36* genes of *Klebsiella* spp. and reference genomes.** (A) BLAST searches of sequence data held at NCBI did not identify any other *Klebsiella* strains carrying a *tnpA* insertion in the 5' end of *ompK35*. However, such an insertion is seen in *E. coli* 1290/03 which has a *tnpA* inserted 45 bp upstream of the start codon of the *ompK35* homolog (*ompF*). A non-identical but similar *tnpA* insertion is seen 42 bp upstream of the start codon of the *ompK36* gene of *K. pneumoniae* WJ19, and 45 bp upstream of the start codon of the *ompK36* homolog (*ompC*) in *E. coli* strains 1290/03, 917/05 and 1566/03. Alternate means disrupting the functionality of the porin encoding genes were also observed with SNPs that generate premature stop codons indicated by red lines. GenBank accession numbers of the sequences are as follows: *E. coli* 1290/03, GQ465829; *K. pneumoniae* WJ19, Q4557043; *E. coli* 917/05, GQ167039; *E. coli* 1566/03, GQ167038. Sequence comparisons were created with ViPTree[84]. (B) OmpK36 amino acid sequences from FK688, other *K. quasipneumoniae* subsp. *similipneumoniae* (Kqs) strains and a *K. pneumoniae* strain (Kpn) extracted from the NCBI database. The conceptual translation of the OmpK36 encoded by the Kpn KP549/04 strain[85] had the same 16-amino acid deletion as FK688 (blue box). Residues showing 100% identity among sequences are highlighted in red. Similarity among sequences is indicated by black bold font highlighted in yellow. Accession numbers for the proteins shown are as follows: UNMC_7493: OVT66776.1; G1129: OVX13856.1; KP-24175: KYZ7217.4; CAB1577: OVX38519.1; KPPSTH03: OYM42369.1; KPCTRSRTH03: TNJ78288.1; ATCC 700603: AWO62949.1; MGH44: ESM62951.1; SWT10: TWV32269.1; KP549/04: ADG56566.1; SB610: VGP86012.1; G747: AZJ27052.1. Sequence alignments were performed with Clustal Omega[86]and rendered with ESPript 3.0[87].

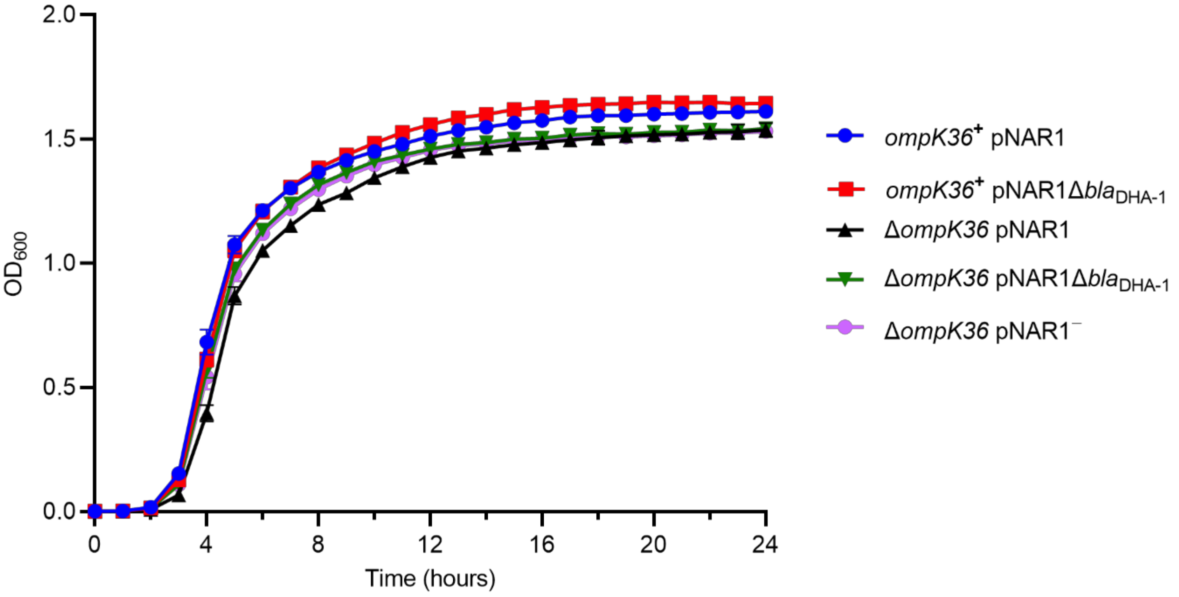


**Figure S4. Growth rate analysis of *K. quasipneumoniae* strains.** Bacteria were cultured in CAMHB and the $OD_{600}$ was measured every hour for 24 hours. Error bars represent SD (n=3). The genotypes of the strains are indicated in the legend. Note that the parental FK688 strain is Δ*ompK36* pNAR1. Where indicated, the other isogenic strains express a functional OmpK36 porin (i.e. *ompK36*$^{+}$), carry the complete pNAR1 plasmid or pNAR1 with a 17 kb (*tnpA-sul1*) deletion (pNAR1*Δbla*$_{DHA-1}$) or have been cured of the plasmid (pNAR1$^{-}$).

A

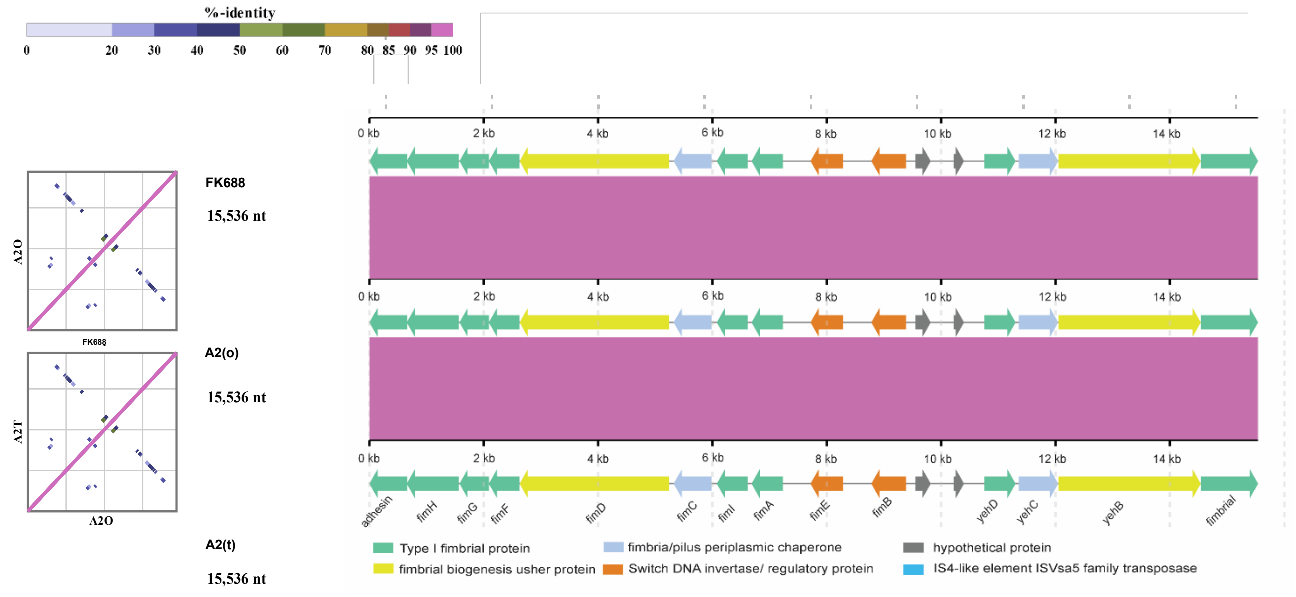


B

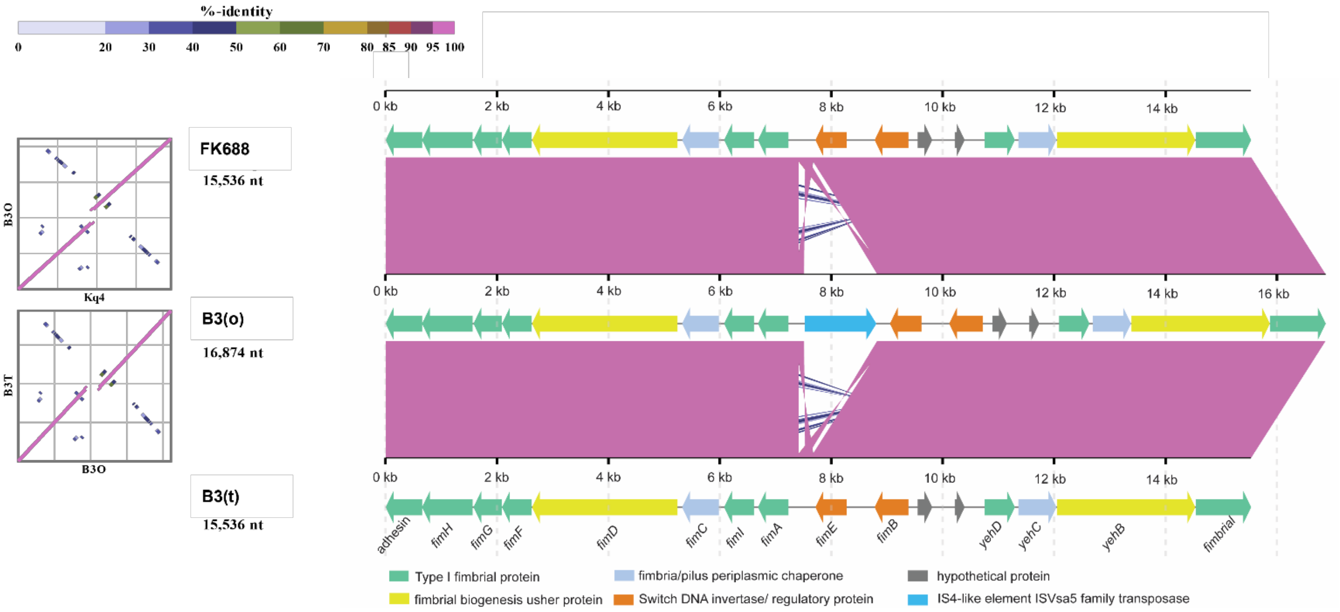


**Figure S5. Comparative sequence analysis of the *fim* gene cluster in evolved *K. quasipneumoniae* strains.** (A) Schematic of the FK688 and its evolved strains A2(o) and A2(t) *fim* gene cluster with 100 % gene similarity between the strains. (B) Sequence comparison of FK688 *ompK36*$^{+}$ revealed a tnpA gene from the IS*4* family (blue arrow) inserted upstream of *fimE* in B3(o). Sequence comparisons were performed with ViPTree[84] .

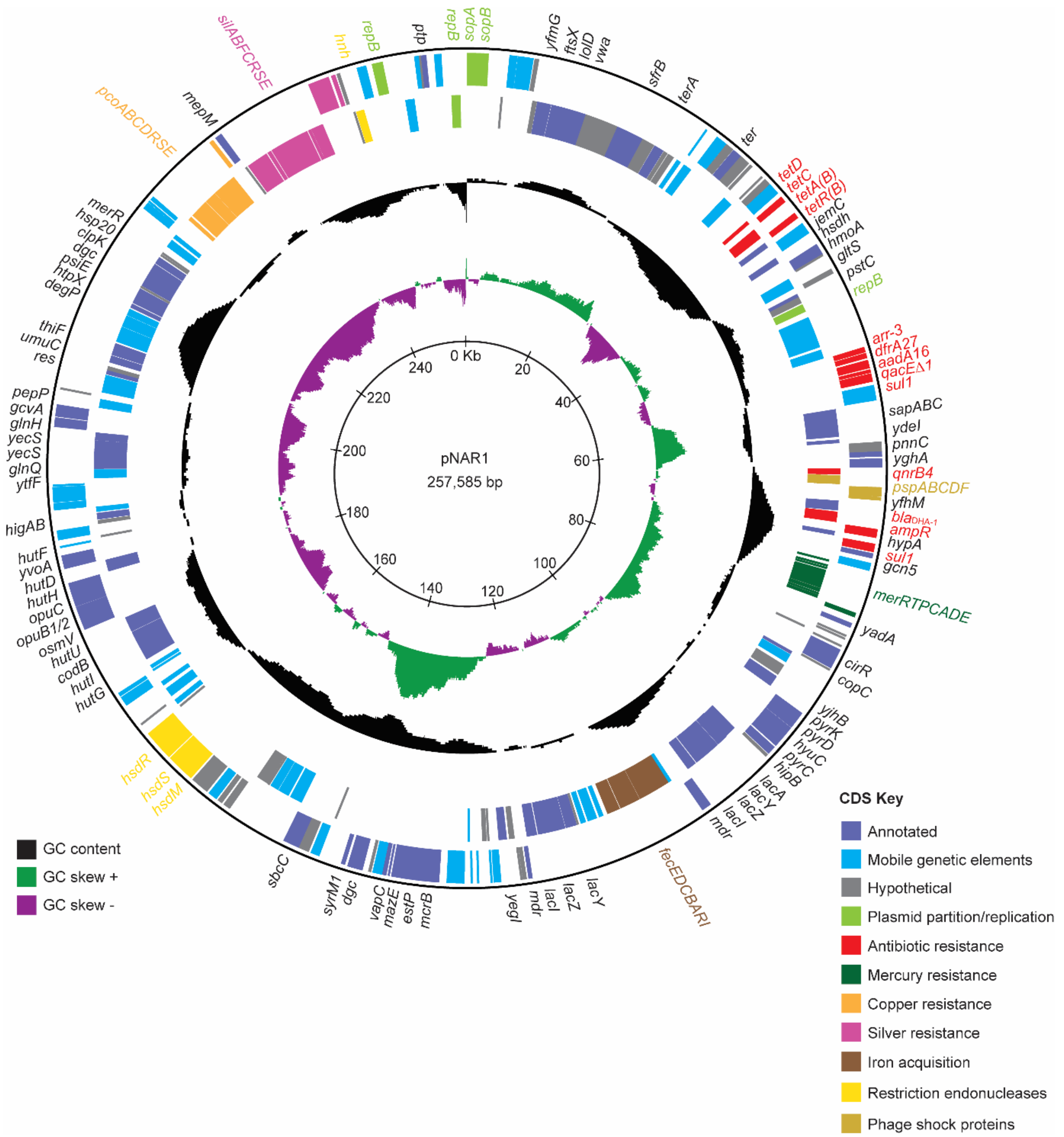


**Figure S6. Detailed physical map of the FK688 plasmid pNAR1.** The coloured lines in the outer concentric circles represent the location of predicted coding sequences in the forward and reverse DNA strands. Annotated CDS are shown in dark blue and labelled with gene names. Hypothetical genes of unknown function are in grey. Mobile genetic elements are in light blue. The position of genes encoding antibiotic resistance determinants (red), mercury (*mer*) resistance (green), copper (*pco*) resistance (orange), and silver (*sil*) resistance (pink) are indicated. The genes encoding a Fe (3+) dicitrate ABC transporter system are in dark brown. The Type 1 *hsdRMS* restriction-modification and H-N-H endonuclease (*hnh*) systems (yellow) and phage shock protein (Psp) system (light brown) are indicated. The middle circle (black) indicates the % GC content, and the inner circle indicates the positive (green) and negative (purple) GC skew [(G-C)/(G+C)]. The map was generated with DNAPlotter [79].

**Supplementary Table S1. Antibiotic resistance genes identified in pNAR1**

| ID | Protein | Gene | Resistance Mechanism | Antimicrobial class |
|---|---|---|---|---|
| 036 | Transcriptional activator of SoxS/MarA/Rob regulon genes | *tetD* | Antibiotic efflux (regulation) | Multiple |
| 037 | Transcriptional repressor of *tetCD* | *tetC* | Antibiotic efflux (regulation) | Multiple |
| 038 | Tetracycline efflux MFS transporter | *tetA*(B) | Antibiotic efflux | Tetracycline |
| 039 | Transcriptional repressor of *tetA*(B) | *tetR*(B) | Antibiotic efflux (regulation) | Tetracycline |
| 054 | Rifampin ADP-ribosyl transferase | *arr-3* | Antibiotic inactivation | Rifamycin |
| 055 | Dihydrofolate reductase | *dfrA27* | Antibiotic target replacement | Diaminopyrimidine |
| 056 | Aminoglycoside 3"-O-nucleotidyltransferase | *aadA16* | Antibiotic inactivation | Streptomycin |
| 057 | Quaternary ammonium compound efflux SMR transporter | *qacΔE1* | Antibiotic efflux | Macrolide |
| 058 | Dihydropteroate synthase | *sul1* | Antibiotic target replacement | Sulfonamide |
| 068 | Quinolone resistance pentapeptide repeat protein | *qnrB4* | Antibiotic target protection | Fluoroquinolone |
| 076 | Class C beta-lactamase DHA-1 | $bla_{\text{DHA-1}}$ | Antibiotic inactivation | Cephamycin, Cephalosporin |
| 077 | Transcriptional repressor of *ampC* | *ampR* | Antibiotic inactivation (regulation) | Cephamycin, Cephalosporin |
| 079 | Dihydropteroate synthase | *sul1* | Antibiotic target replacement | Sulfonamide |

### Supplementary Table S2. Transmembrane transporter systems identified in pNAR1

| ID | Transporter system | Gene/operon | Predicted function |
|---|---|---|---|
| 010 | ABC transporter permease (FtsX) | *salY* | Antimicrobial peptide transport system |
| 011 | ABC transporter ATP-binding protein (LolD) | *lolD* | Lipoprotein export |
| 060-062 | Peptide ABC transporter system (SapABC) | *sapABC* | Antimicrobial peptide transport system |
| 084-090 | Mercury resistance system | *merR, merTPCADE,* | Hg(2+) reduction and inactivation |
| 101 | MFS transporter permease (YjhB) | *araJ* | Arabinose efflux |
| 110, 126 | MFS transporter lactose permease (LacY) | *lacY* | β-galactoside transport |
| 115-118 | Fe(3+) dicitrate ABC transporter system (FecEDCB) | *fecEDCB* | Fe(3+) translocation |
| 189-192 | ABC osmoregulated transporter system (OsmV/OpuBC) | *osmV/opuBC* | Proline/glycine betaine transport |
| 209 | DMT family transporter (FtfF) | *rhaT* | Drug/metabolite transport permease |
| 210-213 | ABC amino acid transport system (GlnQ/YecS/GlnH) | *glnQ/yecS/glnH* | Polar amino acid import |
| 245-251 | Copper resistance system | *pcoABCDRSE* | Sequestration and export |
| 257-263 | Silver resistance system | *silRSE, silCFBAP* | Sequestration and efflux RND transport |

**Supplementary Table S3. β-lactamase prediction and classification using DeepBL.**

| Score [a] | Class [b] | Gene | BLDB BLAST | CDART classification |
|---|---|---|---|---|
| 0.999913 | C | *bla*$_{\mathrm{DHA-1}}$ | DHA-1 β-lactamase | Transpeptidase family |
| 0.999877 | A | *bla*$_{\mathrm{OKP-B-21}}$ | OKP-B-21 β-lactamase | Transpeptidase family |
| 0.997564 | C | *bla*$_{\mathrm{SHK}}$ | SEC-2 SST-1 Serine hydrolase | Transpeptidase family |
| 0.996537 | B | *bla*$_{\mathrm{OPHC}}$ | L1-11 L1 β-lactamase, MBL fold metallo-hydrolase | MBL fold metallo-hydrolase |
| 0.992373 | A | *bla*$_{\mathrm{DAE}}$ | SHV-105 extended-spectrum β-lactamase, D-alanyl-D-alanine endopeptidase | Transpeptidase family |
| 0.989845 | B | *bla*$_{\mathrm{TRN}}$ | OXA-657; OXA-23 family carbapenem-hydrolyzing class D β-lactamase OXA657 | AlaX tRNA deacylase |
| 0.989458 | C | *bla*$_{\mathrm{ABH}}$ | OXA298 hypothetical protein, Alpha/β-hydrolase | Alpha/beta hydrolase |
| 0.974344 | A | *bla*$_{\mathrm{DAC}}$ | SGM-5 β-lactamase class A, Serine-type D-Ala-D-Ala carboxypeptidase | Transpeptidase family |

[a] For each gene, a five-dimensional vector was generated, representing the scores of classes of non-β-lactamase, Class-A, Class-B, Class-C, and Class-D.

[b] The class corresponding to the largest score was treated as the final predicted class for the gene

### Supplementary Table S4. List of plasmids used in this study

| Plasmid | Relevant characteristics[1] | Source/reference |
|---|---|---|
| pKD4 | Contains kanamycin resistance cassette (*kan*) flanked by FRT sites (FRT-*kan*-FRT); *oriR6K*, $Amp^R$, $Km^R$ | (Datsenko & Wanner, 2000)[76] |
| pJET1.2/blunt | Blunt-end cloning vector for insertion of DNA fragments with single deoxyadenosine overhangs; $Amp^R$ | Thermo Scientific™ |
| pDonor(OmpK36) | pJET1.2/blunt carrying FRT-*kan*-FRT and *K. quasipneumoniae* FK688 *ompK36* regions (donor plasmid for lambda Red recombination-mediated repair of *ompK36* gene in FK688); $Amp^R$, $Km^R$ | This study |
| pACBSR | Arabinose-inducible promoter; I-*Sce*I endonuclease; λ-Red recombination genes, $Cm^R$ | (Herring, Glasner & Blattner, 2003)[66] |
| pFLP-BSR | pACBSR carrying fragment length polymorphism (FLP) recombinase to excise the kanamycin cassette, in place of I-*Sce*I; $Cm^R$ | (Rocker *et al.*, 2020)[29] |
| pJP-CmR | Derivative of pJP168 for anhydrotetracycline inducible protein expression. $Cm^R$ | (Rocker *et al.*, 2020)[29] |
| pJP-$bla_{DHA\text{-}1}$ | pJP-Cm containing *ampC* from FK688 | This study |
| pJP-$bla_{OKP\text{-}B\text{-}21}$ | pJP-Cm containing *bla* from FK688 | This study |
| pJP-$bla_{SHK}$ | pJP-Cm containing *CKCOFDID_01495* from FK688 | This study |
| pJP-$bla_{OPHC}$ | pJP-Cm containing *CKCOFDID_02113* from FK688 | This study |
| pJP-$bla_{DAE}$ | pJP-Cm containing *pbpG* from FK688 | This study |
| pJP-$bla_{TRN}$ | pJP-Cm containing *CKCOFDID_04153* from FK688 | This study |
| pJP-$bla_{ABH}$ | pJP-Cm containing *dhmA* from FK688 | This study |
| pJP-$bla_{DAC}$ | pJP-Cm containing *dacB* from FK688 | This study |

[1] Amp, ampicillin; Km, kanamycin; Cm, chloramphenicol.

**Supplementary Table S5. MIC analysis of *E. coli* BW25113 expressing candidate β-lactamases.**

| Antimicrobial drug | MIC (µg/mL) [a] | | | | | | | | | |
|---|---|---|---|---|---|---|---|---|---|---|
| | pJP-CmR | No Vector | pJP-*bla*$_{DHA-1}$ | pJP-*bla*$_{OKP-B-21}$ | pJP-*bla*$_{SHK}$ | pJP-*bla*$_{OPHC}$ | pJP-*bla*$_{DAE}$ | pJP-*bla*$_{ABH}$ | pJP-*bla*$_{DAC}$ | pJP-*bla*$_{TRN}$ |
| **Cephems** | | | | | | | | | | |
| Cefazolin | 2 | 2 | 256 | 8 | 2 | 2 | 1 | 2 | 2 | 2 |
| Cefotaxime | 0.06 | 0.06 | 32 | 0.06 | 0.03 | 0.03 | 0.03 | 0.06 | 0.06 | 0.03 |
| Ceftazidime | 0.25 | 0.25 | 128 | 1 | 0.12 | 0.06 | 0.06 | 0.12 | 0.25 | 0.25 |
| **Carbapenems** | | | | | | | | | | |
| Ertapenem | 0.016 | 0.016 | 0.03 | 0.008 | 0.008 | 0.008 | 0.008 | 0.008 | 0.008 | 0.008 |
| Imipenem | 0.25 | 0.25 | 0.5 | 0.12 | 0.25 | 0.12 | 0.12 | 0.12 | 0.12 | 0.25 |
| Meropenem | 0.016 | 0.016 | 0.03 | 0.016 | 0.016 | 0.016 | 0.016 | 0.016 | 0.016 | 0.016 |

[a] Drug-sensitive, black text; drug-resistant, red-text

**Supplementary Table S6. MIC analysis of pNAR1Δ*bla*$_{DHA-1}$ strains expressing DHA-1**

| Antimicrobial drug | MIC (µg/mL) [a] | | | | | |
|---|---|---|---|---|---|---|
| | **Δ*ompK36* pNAR1Δ*bla*$_{DHA-1}$** | | **Δ*ompK36* pNAR1$^{-}$** | | ***ompK36*$^{+}$ pNAR1Δ*bla*$_{DHA-1}$** | |
| | **pJP-CmR** | **pJP-*bla*$_{DHA-1}$** | **pJP-Cm** | **pJP-*bla*$_{DHA-1}$** | **pJP-Cm** | **pJP-*bla*$_{DHA-1}$** |
| Ceftazidime | 0.5 | 1024 | 0.5 | 1024 | 0.5 | 512 |
| Ertapenem | 0.5 | 32 | 1 | 16 | 0.016 | 0.5 |
| Imipenem | 0.25 | 4 | 0.25 | 4 | 0.12 | 0.25 |
| Meropenem | 0.12 | 2 | 0.12 | 1 | 0.016 | 0.03 |

[a] Drug-sensitive, black text; drug-resistant, red-text

### Supplementary Table S7. FK688 OmpK36$^{+}$, B3(o) and B3(t) genome modification and SNP analysis

| Location | FK688 (Ompk36$^{+}$) | B3(o) | B3(t) | Gene/locus tag | Annotation | Comments |
|---|---|---|---|---|---|---|
| **SNP** | | | | | | |
| Chromosome | A | C | C | *smc_1* | Chromosome partition protein Smc/Phage tail fiber protein | multiple aa replacement (all strains 4195 aa) |
| Chromosome | G | A | A | *LKCPGFBJ_02921* | Phage tail assembly | aa replacement alanine (A) --> valine (V) in B3O and B3T |
| Plasmid | T | C | C | *kdgK_2* | IS3 family element, transposase | Frameshift mutation in B3O and B3T |
| Plasmid | N | C | C | *kdgK_2* | IS3 family element, transposase | Frameshift mutation in B3O and B3T |

| Location | FK688 (Ompk36$^{+}$) | B3(o) | B3(t) | Product | Gene(s) affected |
|---|---|---|---|---|---|
| **Gene modification** | | | | | |
| Chromosome | 473 aa | 61 aa/ 398aa | 473 aa | Deletion of 14 aa = 2 fractions of the protein in B3O | PEP-dependent dihydroxyacetone kinase, phosphoryl donor subunit DhaM |
| Chromosome | N/A | Insertion | N/A | IS4-like element ISVsa5 family transposase | Intergenic: Fimbrial subunit type 1 (*fimA*) and Type 1 fimbrial regulatory protein *fimE* |
| Chromosome | N/A | N/A | Insertion | IS4-like element ISVsa5 family transposase | Intergenic: Type 3 fimbria minor subunit MrkF and phosphodiesterase MrkJ |
| Chromosome | N/A | N/A | Insertion | IS4-like element ISVsa5 family transposase | Intragenic: UDP-glucose: undecaprenyl-phosphateglucose-1-phosphate transferase (*wcaJ*) |
| Chromosome | N/A | N/A | Insertion | IS4-like element ISVsa5 family transposase | Intergenic: IS4-like element ISVsa5 family transposase and Putative ATP-binding component of a transport system |
| Chromosome | N/A | N/A | Insertion | IS4-like element ISVsa5 family transposase | Intragenic: ABC transporter, substrate-binding protein KPN |

**Supplementary Table S8. FK688, A2(o) and A2(t) genome modification and SNP analysis**

| Location | FK688 | A2(o) | A2(t) | Gene | Annotation | Comments |
|---|---|---|---|---|---|---|
| **SNP** | | | | | | |
| Chromosome | G | C | C | *LptB_3* | LPS export system ATP binding protein LptB | 2 ORF LptB in FK688 |
| Chromosome | T | C | C | *LptB_3* | LPS export system ATP binding protein LptB | 2 ORF LptB in FK688 |
| Chromosome | A | C | C | *WcaA* | colanic acid biosynthesis glycosyl transferase WcaA | No changes in protein alignment |
| Chromosome | A | A | G | *novR* | Decarboxylase NovR | change lysine (K) --> glutamic acid (E) in A2T |
| Chromosome | A | G | G | *smc_1* | Chromosome partition protein Smc. | 4126 aa in Kq1 --> 4195 aa in A2O & A2T |
| Chromosome | A | C | C | *smc_1* | Chromosome partition protein Smc. | 4126 aa in Kq1 --> 4195 aa in A2O & A2T |
| Chromosome | G | A | A | *smc_1* | Chromosome partition protein Smc. | 4126 aa in Kq1 --> 4195 aa in A2O & A2T |
| Chromosome | A | C | C | *smc_1* | Chromosome partition protein Smc. | 4126 aa in Kq1 --> 4195 aa in A2O & A2T |
| Chromosome | A | G | G | *smc_1* | Chromosome partition protein Smc. | 4126 aa in Kq1 --> 4195 aa in A2O & A2T |
| Chromosome | A | G | G | *smc_1* | Chromosome partition protein Smc. | 4126 aa in Kq1 --> 4195 aa in A2O & A2T |
| Chromosome | C | T | T | *smc_1* | Chromosome partition protein Smc. | 4126 aa in Kq1 --> 4195 aa in A2O & A2T |
| Chromosome | A | G | G | *smc_1* | Chromosome partition protein Smc. | 4126 aa in Kq1 --> 4195 aa in A2O & A2T |
| Chromosome | G | T | G | *cad* | Transcriptional activator of cad operon | change asparagine (N) --> lysine (K) in A2O |
| Plasmid | A | G | G | *kdgK_2* | IS3 family element, transposase | Frameshift mutation leucine(L) --> proline (P) in A2O and A2T |
| Plasmid | T | C | C | *kdgK_2* | IS3 family element, transposase | Frameshift mutation in A2O and A2T |
| Plasmid | N | C | C | *kdgK_2* | IS3 family element, transposase | Frameshift mutation in A2O and A2T |
| **Location** | **FK688** | **A2(o)** | **A2(t)** | **Type** | **Product** | **Gene(s) affected** |
| **Gene modification** | | | | | | |
| Chromosome | N/A | N/A | Insertion | Insertion | IS4-like element ISVsa5 family transposase | Within the gene: N,N'-diacetylchitobiose-specific 6-phospho-beta-glucosidase (*chbF*) |
| Chromosome | N/A | N/A | Change | Change | aa replacement aspartic acid (D) for glutamic acid (E) | outer membrane porine_Maltoporin _LamB_3 |

### Supplementary Table S9. List of strains used in this study

| Strain | Relevant characteristics[1] | Source or reference |
|---|---|---|
| ***K. quasipneumoniae*** | | |
| FK688 $\Delta$*ompK36* pNAR1 | Wildtype, clinical isolate from a bloodstream infection case from the First Affiliated Hospital of Wenzhou Medical University, China. Expresses beta-lactamase $bla_{OKP\text{-}B\text{-}21}$. Deficient in *ompK35* and *ompK36* porin expression. Harbours a 258 kb plasmid pNAR1 ($Amp^R$, $Tet^R$, $Rif^R$, $Trp^R$, $Stp^R$, $Ery^R$, $Sdz^R$, $Cip^R$). | (Bi et al., 2017)[23] |
| $\Delta$*ompK36* pNAR1$\Delta bla_{DHA\text{-}1}$ | FK688 with a 17 kbp deletion from *tnpA-sul1* in pNAR1. | This study |
| $\Delta$*ompK36* pNAR1$^{-}$ | FK688 cured of pNAR1. | This study |
| $ompK36^{+}$ pNAR1 | FK688 with a genetically repaired and functional *ompK36* gene. Carries pNAR1. | This study |
| $ompK36^{+}$ pNAR1$\Delta bla_{DHA\text{-}1}$ | FK688 with a genetically repaired and functional *ompK36* gene. It has a 17 kb deletion from *tnpA-sul1* in pNAR1. | This study |
| FK688-GFP$^{+}$ | FK688 with a constitutively expressed green fluorescent protein (GFP). GFP gene inserted downstream of the *glmS* gene via pGRG-eGFP. | This study |
| A2(o) $\Delta$*ompK36* pNAR1$\Delta bla_{DHA\text{-}1}$ | Evolved strain from Kq1. It has a 17 kb deletion from *tnpA-sul1* in pNAR1. Forms opaque colonies. | This study |
| A2(t) $\Delta$*ompK36* pNAR1$\Delta bla_{DHA\text{-}1}$ | Evolved strain from Kq1. It has a 17 kb deletion from *tnpA-sul1* in pNAR1. Forms translucent colonies. | This study |
| B3(o) $ompK36^{+}$ pNAR1$\Delta bla_{DHA\text{-}1}$ | Evolved strain from Kq4. It has a 17 kb deletion from *tnpA-sul1* in pNAR1. Forms opaque colonies. | This study |
| B3(t) $ompK36^{+}$ pNAR1$\Delta bla_{DHA\text{-}1}$ | Evolved strain from Kq4. It has a 17 kb deletion from *tnpA-sul1* in pNAR1. Forms translucent colonies. | This study |
| ***K. pneumoniae*** | | |
| B5055 | Hypermucoviscous phenotype. Wild-type, clinical isolate, serotype K2;O1 | Statens Serum Institut, Denmark |
| B5055 nm | B5055 deletion mutant Δwza-wzc::km (non-mucoid); $Km^R$ | Prof. Richard Strugnell, University of Melbourne |
| AJ097 | Wild-type, clinical isolate, serotype K2; $Amp^R$. Origin: Urine isolate from urinary tract infection case, Melbourne Australia, 2002 | (Jenney, A. W. *et al.*, 2006)[77] |
| ***E. coli*** | | |
| DH5α | $F^{-}$ endA1 *hsd*R17($r_K^{-}$, $m_K^{+}$) *sup*E44 λ– *thi*-1 *rec*A1 *gyr*A96 *rel*A1 *deoR* Δ(*lac*ZYA-*arg*F) U169 Φ80*dlac*ZΔM15; $Nal^R$ | Invitrogen |
| ATCC® 25922™ | CLSI control strain for antimicrobial susceptibility testing | ATCC® |
| BW25113 (WT) | *rrn*B3 Δ*lac*Z4787(::rrnB-3) *hsd*R514 Δ(*araD-araB*)567 Δ(*rhaD-rhaB*)568, *rph*-1 | (Baba *et al.,* 2006)[78] |

[1] Amp, ampicillin; Tet, tetracycline; Rif, rifamycin; Trp, trimethoprim; Stp, streptomycin; Ery, erythromycin; Sdz, sulfadiazine; Cip, ciprofloxacin; Nal, nalidixic acid.

### Supplementary Table S10. List of oligonucleotide primers used in this study

| Primer | Sequence (5-3')[1] | Description |
|---|---|---|
| **Construction of FK688 OmpK36$^{+}$ strains** | | |
| K36_insert-R | gcgcgacctactacttcaacaaaaacatgtccacctatgttgactacaaaatcaacctgctg | Construction of pDonor(OmpK36) plasmid |
| K36_insert-F | gttgaagtagtaggtcgcgcccacgtcaacatatttcaggatgtcctggtcgcc | |
| K36_Km-F | ctaaggaggatattcatatggtcgcaagctgcataacaaa | |
| K36_Km-R | gaagcagctccagcctacacattagaactggtaaaccaggcccag | |
| K36_ISceI-R | tagggataacagggtaatgcccgacggtgatatccatc | |
| K36_ISceI-F | tagggataacagggtaatgcttcggtacctctgtaacttatga | |
| pKD4-F | tgtgtaggctggagctgcttc | Kanamycin cassette from pKD4 |
| pKD4-R | catatgaatatcctccttag | |
| **Cloning of putative β-lactamases genes for anhydrotetracycline-inducible expression** | | |
| blaDHA-1_For_NR | gtccCCATGGtgaaaaaatcgttatctgcaac | |
| blaDHA-1_Rev_NR | cgtcAAGCTTattccagtgcactcaaa | |
| blaOKP_F_NR | tagcGAATTCatgcgttatgttcgcctgtgcc | |
| blaOKP_R_NR | gcatAAGCTTctagcgctgccagtg | |
| blaSHK1_F_NR | gttcCCATGGtgataagaaaaccactggcc | |
| blaSHK1_R_NR | atgcAAGCTTaacgcagctcgcg | |
| blaOPHC2_F_NR | ctagGAATTCatgacaccagctcccttttataccctgac | |
| blaOPHC2_R_NR | acggAAGCTTtcgctgtgatcggtgtt | |
| blaDAE1_F_NR | tgcaGAATTCatgatgccgaaatttcgagtctctttgc | |
| blaDAE1_R_NR | gatcAAGCTTttaatcgttctgcgcg | |
| blaABH1_F_NR | acgtCCATGGTGaacagattatccctgatcc | |
| blaABH1_R_NR | gatcAAGCTTacaaccgatcggcg | |
| blaDAC1_F_NR | aaggCCATGGtgcgatttcccagatttatc | |
| blaDAC1_R_NR | aagcAAGCTTtagttgttctggtacaaatcc | |
| blaTRN1_F_NR | cgtaCCATGGtgactgaacgggtttattacac | |
| blaTRN1_R_NR | aatcAAGCTTacgtcagggaatagctgatc | |
| pJPMCS_For | cctaatttttgttgacactctatcattg | pJP-CmR-gene insert sequencing primers |
| pJPMCS_Rev | gccaggcaaattctgttttatcagaccg | |

[1] Restriction endonuclease recognition sites are capitalised.

**Supplementary Materials and Methods.**

***Genome sequencing and evaluation***

In brief, high molecular weight DNA was prepared as a Nanopore sequencing library, according to the manufacturer's protocols using a ligation sequencing kit (SQK-LSK109, Oxford Nanopore), with minor modifications. The optional shearing step was omitted and all mixing steps for the DNA sample was done by gently flicking the microfuge tube instead of pipetting to further reduce DNA shearing.

DNA repair treatment was carried out using NEBNext FFPE DNA Repair Mix (M6630, New England Biolabs). End repair and A-tailing was performed with NEBNext Ultra II End Repair/dA-tailing Module (E7546, New England Biolabs) and the sample was incubated at 20 °C for 5 min and then 65 °C for 5 min. End-repaired product was purified with 1× Agencourt AMPure XP beads.

Adapters provided in the respective library kits were ligated to the DNA with Quick T4 DNA Ligase (M2200L, New England Biolabs) and samples were incubated at room temperature for 10 min. Purification and loading of adapted libraries on an appropriate flow cell (R9.4.1, FLO-MIN106D, Oxford Nanopore) was completed as per the manufacturer's recommendations and sequenced using the appropriate MinKNOW workflow. The library was base-called using Guppy (ont-guppy-for-gridion, 3.0.6). Reads with a length less than 1,000 bp were discarded.

The sample had an estimated Nanopore coverage of 884-fold, with an average read length of 7,445 bp and a range of 1,000 bp to 191,716 bp.

Illumina sequences were prepared on a NextSeq 500 platform, with 150 bp paired-end chemistry. Reads were trimmed to remove adaptor sequences and low-quality bases with Trimmomatic (https://github.com/timflutre/trimmomatic), with Kraken used to investigate contamination (v0.10.5-beta, https://github.com/DerrickWood/kraken).

Assembly involved long-read-only assembly of long reads, followed by short-read correction. In brief, Nanopore reads were downsampled using Filtlong (v0.2.0, https://github.com/rrwick/Filtlong) to retain the highest quality reads (10% of all, equivalent to an estimated 88-fold coverage). These reads were assembled using Canu (1.8, https://github.com/marbl/canu), with an expected genome size of 5,000,000 bp. The assembled contigs output by Canu were circularised where appropriate and validated through a read mapping approach in Geneious Prime (2019.2.1), before short-read correction. The corrected assembly was oriented to *dnaA* and annotated using Prokka (https://github.com/tseemann/prokka).

The sample had varied coverage across assembled molecules, ranging in Illumina coverage between 156-fold for the chromosome and 188-fold for the plasmid.

***Membrane purification and isolation***

Overnight cultures of strains were diluted 1:100 in 200 mL CAMHB and grown until mid-log ($OD_{600}$ of 0.5). Cells were harvested by centrifugation (10,000 × g, 10 min, 4 °C) and resuspended in 10 mM Tris-HCl, pH 7.5. The centrifugation was repeated, and cells resuspended in Tris-Sucrose (TS) buffer (10 mM Tris-HCl, pH 7.5, 0.75 M sucrose). Peptidoglycan was degraded using a final concentration of 50 µg/mL lysozyme and host serine proteases were inhibited using a final concentration of 2 mM phenylmethylsulfonyl fluoride. The outer membrane was destabilised for lysis in 2 volumes of 1.65 mM EDTA, pH 7.5. Cells were incubated on ice for 10 min and then lysed using an AVESTIN Emulsiflex-C3 (4 passes at ~ 15,000 psi). Cell lysates were subjected to centrifugation (15,000 × g, 20 min, 4 °C) to remove cell debris. The supernatant was collected, and total membranes were isolated by ultracentrifugation (132,000 × g, 45 min, 4 °C) using a 70.1 Ti rotor. Membrane pellets were resuspended and pooled from duplicate samples in ~ 8 mL TES buffer (2.2 mM Tris-HCl, pH 7.5, 1.1 mM EDTA, 0.25 M sucrose) and subjected to ultracentrifugation as before. Membrane pellets were resuspended in 200 µL of 25% sucrose in 5 mM EDTA, pH 7.5, and stored at -80 °C.

***Protein expression***

For all membrane extract samples, total protein content was quantified with a NanoDrop 1000 Spectrophotometer (Thermo Scientific) and standardised to equivalent concentrations. Samples (~ 2 µg) were loaded onto a sodium dodecyl sulfate-polyacrylamide gel (SDS-PAGE) containing 11% (wt/vol) 37.5:1 acrylamide-bisacrylamide, 0.375 M Tris (pH 8.8), 0.1% (wt/vol) SDS, and 0.5 mM EDTA in the separating gel and 4% (wt/vol) 37.5:1 acrylamaide-bisacrylamide, 0.25 M Tris (pH 6.8), 0.1% (wt/vol) SDS, and 0.5 mM EDTA in the stacking gel. The proteins in the gel were transferred to a 0.2 µm nitrocellulose membrane (Bio-Rad) using a Trans-Blot Turbo Transfer System (Bio-Rad) (0.6 A, 25 V, 15 min), and the nitrocellulose filter was thereafter incubated overnight in TBST (tris-buffered saline; 0.1% Tween 20) containing 5% skim milk.

OmpK36 was detected by western blot using polyclonal antibodies raised in rabbits against the *E. coli* homologue OmpC, which is cross-reactive to OmpK35 and OmpK36 in *Klebsiella*. For use as a loading control, the outer-membrane protein BamD was detected using an αBamD antibody raised in rabbits against the *E. coli* BamD homologue. Goat Anti-Rabbit IgG antibody, HRP-conjugate (Sigma-Aldrich) was used as the secondary antibody. All antibodies were used at a 1:20,000 dilution in TBST containing 2% skim milk. Proteins were detected by chemiluminescence.

***Capsule extraction***

Overnight bacterial cultures were sub-cultured 1:50 in LB media and grown until mid-log phase ($OD_{600}$ ~ 0.4-0.6). Cells were collected by centrifugation (5,000 × *g*, 15 min, 5 °C), and the pellet resuspended in 1 mL of $H_2O$. After centrifugation (14,000 × *g*, 10 min), the pellet was resuspended in 500 µL $H_2O$ and viable counts were determined. Samples were incubated at 68 °C for 2 min and 500 µL phenol (Sigma #P4557) added and mixed by inversion. Following incubation at 68 °C for 30 min the mixture was cooled on ice for 2 min. Next, 500 µL chloroform (Sigma-Aldrich #472476) was added and the mixture inverted. Samples were centrifuged (10,000 × *g*, 5 min) and approximately 400 µL of cell-free supernatant was mixed with 1 mL absolute ethanol and incubated at -20 °C for 20 min and washed with 70% ethanol. The carbohydrate-containing precipitate was resuspended in 500 µL $H_2O$. Samples were stored at 4°C.

***Capsule quantification***

Capsular material (200 µL) was mixed with 1.2 mL of 12.5 mM sodium tetraborate (Borax; $NaBH_4$) in concentrated $H_2SO_4$. The mixtures were vigorously vortexed, boiled for 5 min, and then cooled on ice for 10 min before adding 20 µL of 0.15% 3-hydroxydiphenol in 0.5% NaOH. Absorbance was measured at 520 nm. A duplicate sample from each strain was prepared as described above but treated with 0.5% sodium hydroxide alone and used to measure the background absorbance at 520 nm. The glucuronic acid concentration in background-subtracted values for each sample was determined from a standard curve of D-Glucuronic acid.
